# Host Physiology–Dependent Lysis Timing Shapes Bacteriophage Competition Under Nutrient Fluctuations

**DOI:** 10.1101/2025.11.29.690970

**Authors:** Anja K. Ehrmann, Namiko Mitarai

**Affiliations:** The Niels Bohr Institute, University of Copenhagen, Copenhagen, Denmark

## Abstract

When bacteriophages infect starved host bacteria, the restrictive host physiology may lead to prolonged latent periods and/or reduced burst sizes, compared to infection of a fast-growing bacterium. Using a mathematical model, we explore a system of two types of virulent phages that have distinct host physiology dependencies and are infecting a shared bacterial host population. We consider different environmental regimes to test whether they can compete and coexist under fluctuating conditions, putting emphasis on phases with limited resources for bacterial growth. We find that the fitness of a phage that can modulate lysis timing in response to changes in the host physiology is elevated in fluctuating feast-famine environments compared to more stable environments which favor rapid lysis with a reduced burst size. This effect is closely coupled to the increased mortality of free phages due to abortive adsorption to already infected host bacteria during starvation phases. We identify specific system dynamics that either support or suppress the propagation of the delayed lysis phage. This theoretical analysis highlights the competitive benefits and limitations of lysis delay as a phage propagation strategy. Our results underscore the importance of considering the bacterial physiology dependence of bacteriophage replication in order to correctly predict phage fitness and population dynamics in complex environments.

**IMPORTANCE:** Bacteriophage replication depends strongly on the physiological state of the host, yet most ecological and theoretical studies treat phage life histories as fixed traits. This overlooks how nutrient limitation, starvation, and fluctuating growth conditions reshape infection outcomes. By examining competition between phages with different responses to host physiology, our work shows how environmentally driven changes in the latent period can alter which phages persist, spread, or are lost. These insights clarify when delayed lysis is a beneficial strategy and when it becomes a liability. More broadly, our results highlight the need to integrate host physiology into models of phage–host dynamics to better understand microbial ecosystems and to guide applications such as rational phage therapy design.

## INTRODUCTION

Bacteriophages are gaining increased attention due to their central role in shaping the dynamics of microbial ecosystems and their potential application as therapeutic agents. Considering that the infection dynamics of different types of phages vary widely, even among strictly lytic phages, we can expect to observe complex interactions and environmental dependencies when multiple phages with overlapping host specificity are present in the same system. Understanding phage replication and competition across diverse environmental contexts is essential for predicting microbial ecosystem dynamics.

Phages are effective agents of managing bacterial communities and change growth dynamics, community composition, and metabolic activity of different ecosystems [1, 2, 3]. In a community without phages, and under the assumption of one shared, limiting growth substrate, the fastest growing bacterial species will dominate the population quickly and drive other strains to extinction through competitive exclusion. However, the introduction of phages to the system can alter these dynamics and even reverse them [4, 5, 6, 7]. The phage species infecting the fastest growing strain will replicate quickly and therefore suppress its host bacterium’s growth, opening opportunities for slower growing strains to thrive.

In complex microbial ecosystems, phages must navigate multiple factors that can promote or inhibit successful infection and replication. Phages and bacteria are engaged in a continuous “arms race”, with bacteria developing resistance to infection and phages evolving to circumvent these defenses [8, 9, 10, 11, 12]. Additionally, phages compete against other phages and mobile genetic elements for available hosts, and must “defend” their host cell against superinfection after the successful takeover of a host cell [13, 14].

It is assumed that bacteria spend most of the time under conditions of nutrient limitation in their natural environments [15, 16, 17, 18, 19]. Consequently, phage infections often occur under conditions that differ significantly from the pure cultures and rich growth media that serve as the default experimental model to study phage infection in the laboratory. The parameters that characterize phage replication (adsorption rate, burst size, latent period) are commonly used to define phage fitness, but these parameters can vary significantly depending on the physiological state of the host bacterium. Phage production relies on the infected host’s gene expression machinery, so that slow bacterial growth will lead to slower or even completely halted phage production [20, 21, 22].

Despite this variability, theoretical analyses often treat these traits as constant parameters [4, 5, 6], or assume simple linear relations to the growth rate in cases where the growth physiology dependence is important for the system [23, 24, 25]. While some studies have characterized how phage replication parameters change with host physiology for selected model phages [20, 21, 26, 27], and others have documented the stochastic variability of these parameters [28, 29, 22], broader datasets on phage infection dynamics under suboptimal conditions are lacking. Similarly, the consequences of different host physiology dependencies for community dynamics remain underexplored. Even though the available data is limited, remarkably different responses to infection of starved *Escherichia coli* cells have been observed. For instance, phage T4 is capable of delaying lysis when infecting a non-growing starved cell. In this situation, T4 initiates the early infection steps but halts replication before assembling progeny [30]. The viral life cycle resumes when nutrients are restored, even after long starvation periods. Note that this observation is a separate phenomenon from the T-even phages’ lysis inhibition triggered by the adsorption of multiple phages to the same host [31, 32], though the phenotypic consequence, delayed lysis, is similar. Other phages, e.g. T7, maintain a strictly lytic strategy, continuing to replicate and lyse host cells even during starvation [33, 34]. The burst size of T7 has been shown to be dependent on the host growth rate under steady-state growth (similar to T4) [21]. However, the extent to which T7’s burst size is reduced when a previously fast-growing host population enters stationary phase/starvation remains an open question [21, 34]. The ability to produce phage progeny from starved hosts has also been observed for other phages, e.g. phages UT1 and ACQ of *Pseudomonas aeruginosa* [34]. Conversely, infection of a stationary phase host is detrimental to virulent mutants of phage *λ*, with no functional progeny produced [22].

It is logical that phages would react with either reduced burst size or longer latent periods to diminished host physiology. When the host metabolism is limited to a degree that is unable to support the usual phage production rate, a lytic phage can either ‘hibernate’ and wait for conditions that enable lysis with a full burst size or lyse after the usual latent period with a smaller burst size. Note that this separation into different lysis strategies under host starvation is only indirectly related to the work that has been done to find optimal lysis timing for different phages on exponentially growing hosts. If host resources are not limited, phages can in principle continue to accumulate phage progeny many times their usual burst sizes if lysis is not possible due to, e.g., a mutation [35], suggesting that the actual combination of latent period and burst size is the result of an evolutionary optimization process [36, 37, 38, 39, 40, 41, 42].

In this study, we draw inspiration from the replication strategies of phage T4 (hibernation) and phage T7 (continuous bursting) to model two otherwise identical virulent phages in direct competition for a single host population experiencing cycles of abundant and limited resources (feast-famine cycles). These two approaches represent fundamentally different life-history strategies for coping with resource limitation. By modeling two abstract phages only loosely inspired by replication strategies observed in natural phages, we try to isolate the specific system dynamics and prerequisites that support the competitive advantage of a delayed lysis strategy.

The delayed lysis strategy we consider is one form of “pseudolysogeny” where infected cells neither proceed towards the lysis pathway nor replicate with the host [43, 44]. This strategy may provide better survival of phages in environments harsh to free phage particles, similar to the advantage of lysogenization [45, 46, 47]. On the other hand, continuous lysis enables the phages to spread even in a starved population, as long as enough progeny is produced even in the diminished lysis, as demonstrated in the continuous enlargement of T7 phage plaque in a stationary lawn [48]. Such a behavior will allow the phage to keep relatively high number of free phage particles in the system, allowing them to be ready to infect newly available cells all the time, and to continue diffusing in space, searching for other available hosts.

Yet, it remains unclear which strategy confers a competitive advantage when phages compete for hosts under recurrent nutrient fluctuations, inducing variability in host physiology. Understanding this competition is crucial for predicting phage dynamics in natural ecosystems and for designing robust phage-based therapeutic interventions in complex environments.

## MODEL

### Substrate dependent host growth and two types of phages

We present a mathematical model for two virulent phages infecting a population of susceptible host bacteria in a well-mixed environment. In the first version of the model, we simulate a chemostat culture with constant influx of substrate for bacterial growth and constant dilution of the culture at the same flow rate. In the second model, we simulate repeating cycles of batch cultures, resulting in alternating conditions of abundant growth substrate (feast) and starvation conditions (famine). We model the growth of the uninfected bacterial population *B* under the assumption that the growth rate follows Monod’s growth law [49] as

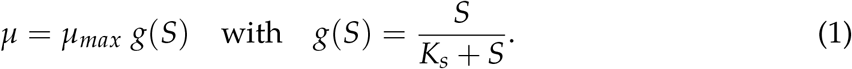

Here *µ*_*max*_ is the maximum growth rate when the nutrient availability is unlimited, and *K*_*s*_ is the half-saturation constant. One unit of the growth substrate *S* is defined as the amount needed to support the duplication of one bacterial cell. In order to contrast the two phage strategies, we consider two types of virulent bacteriophages that are modeled with identical characteristics, except for how their lysis behavior depends on the bacterial growth state.

### Delayed lysis phage

We call one of the phages DL (delayed lysis) and assume that the lysis timing slows down as the bacterial doubling time increases, while the burst size remains constant. More precisely, the latent period of the host cell infected by a DL phage is given by *τ*_*DL*_(*S*) as follows:

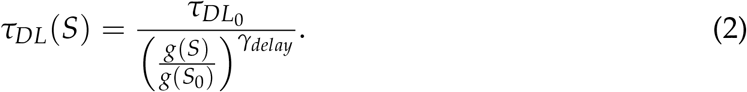

The growth rate *g*(*S*) is normalized to the maximum growth rate *g*(*S*_0_) to ensure that both phages have an identical minimum latent period at the maximum substrate concentration at the start of each feast cycle, or if the maximum substrate concentration is reached in the chemostat system (*τ*_*DL*_(*S*_0_) = *τ*_*RB*_). The parameter *γ*_*delay*_ controls the strength of the lysis delay response. *γ*_*delay*_ = 1 assumes the direct proportionality, as previously proposed in [24], and we use this as a default value. If *γ*_*delay*_ > 1, the latent period of phage DL will increase more rapidly when the bacterial growth rate is decreasing. If *γ*_*delay*_ < 1, phage DL will only slowly increase the latent period in response to the diminishing host physiology.

### Reduced burst size phage

In contrast, the latent period of phage RB (reduced burst) *τ*_*RB*_ is independent of the bacterial growth and remains constant. In order to modulate the competitive strength of phage RB relative to phage DL, we couple the burst size to the substrate concentration through *g*(*S*) with a step-function in the following manner:

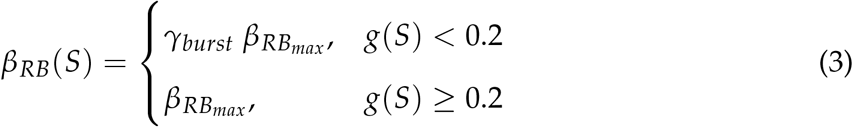

Here, the parameter *γ*_*burst*_ < 1 defines the burst size reduction factor when the growth rate of the host is slow enough. The threshold nutrient concentration *S*_*th*_ for the reduction of phage RB’s burst size is defined by *g*(*S*_*th*_) = 0.2, resulting in *S*_*th*_ = *K*_*s*_/4. The characteristic lysis behavior of the two phages is visualized in Figure **1**.

**FIG 1:**
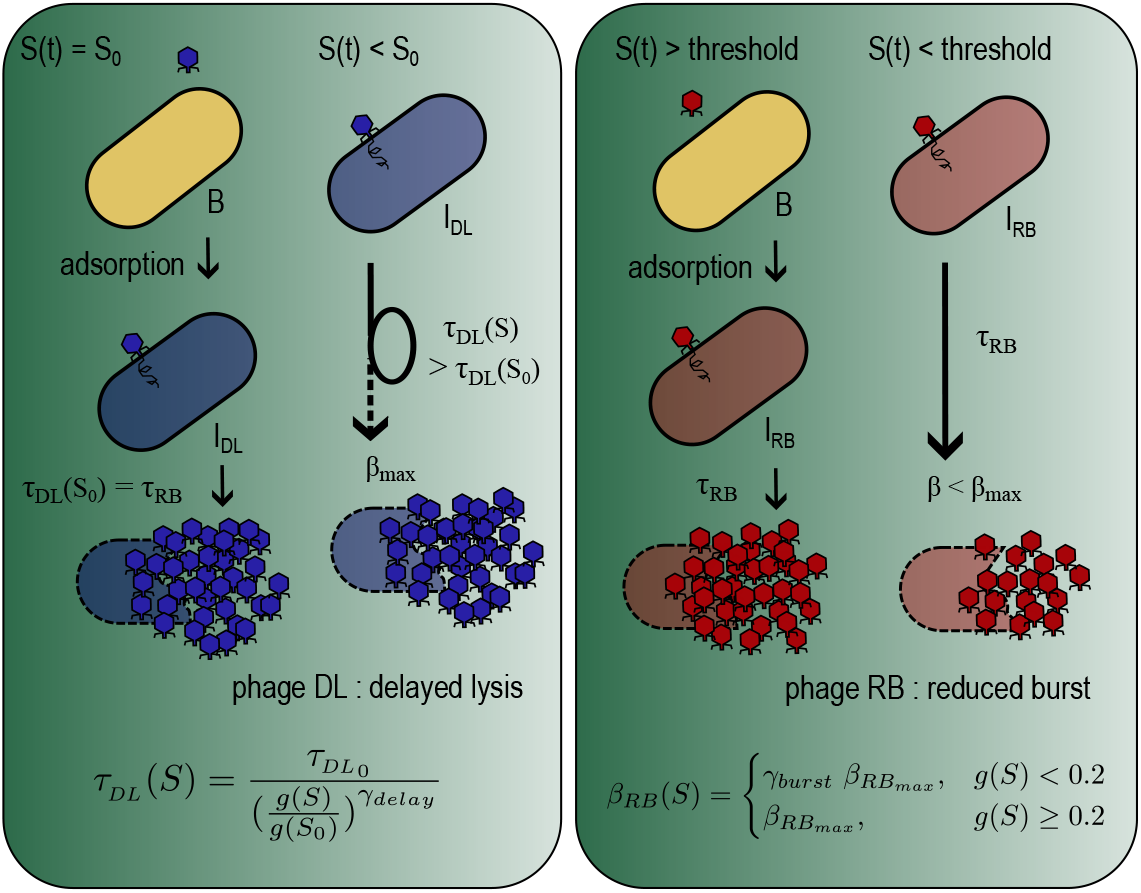
Visual representation of the replication properties of the two phages in the model. The latent period of phage DL (left panel) depends on the host physiology according to eq. 2 so that lysis is delayed relative to phage RB whenever *S*(*t*) < *S*_0_. When the infected cells lyse, they lyse with the full burst size *β*_*max*_. The latent period of phage RB (right panel) is constant, but the burst size is coupled to the host physiology. If *S*(*t*), and therefore the host growth rate, drops below a threshold, the burst size is reduced to *γ*_*burst*_*β*_*max*_ < *β*_*max*_ (eq. 3).

### Model 1 - Chemostat

We consider a well-mixed environment with a constant influx of growth substrate *S* and a constant dilution of the culture at rate *ω*. Within the system we consider the concentration of the bacterial population *B*, the free RB (DL) phage particles *P*_*RB*_ (*P*_*DL*_), the bacteria infected by either RB or DL phage *I*_*RB*_ (*I*_*DL*_) and the concentration of the growth substrate *S*. We assume that the infection of host bacteria follows a Lotka-Volterra type interaction and that the free phage particles adsorb to both uninfected and already infected cells with the adsorption rate *η*. In order to make the latent period distribution to peak around its mean, we model the progression between the primary RB (DL) phage adsorption to the burst as a *n*-step sequential reaction, where the *k*-th state denoted as 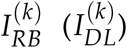, with the identical transition rates to the next steps *n*/*τ*_*RB*_ (*n*/*τ*_*DL*(*S*)_). When an already infected cell adsorbs a phage, the phage particle is inactivated and the status of the cell does not change; in other words, the first infection determines the eventual fate of the infected cell. This is particularly important for phage DL which may experience a long latent period, and is consistent with the T-even phages’ superinfection exclusion mechanism, where DNA injected by secondary adsorbing phages gets trapped and degraded in the periplasm [50, 51, 52]. A bacterium primarily infected by RB (DL) phage will produce *β*_*RB*_(*S*) (*β*_*DL*_) new RB (DL) phage particles that will be released after the latent period of *τ*_*RB*_ (*τ*_*DL*_(*S*)), respectively. Functional forms of *β*_*RB*_(*S*) and *τ*_*DL*_(*S*) are described in eqs. 2 and 3. The full chemostat model is given as follows:

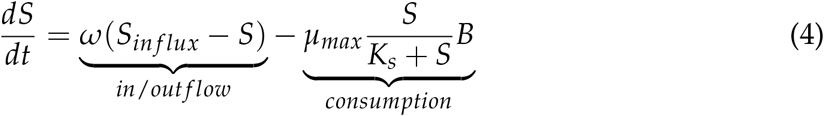

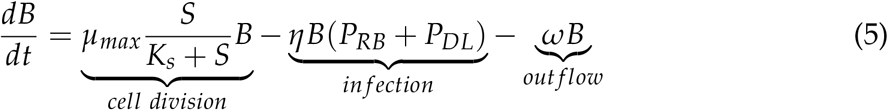

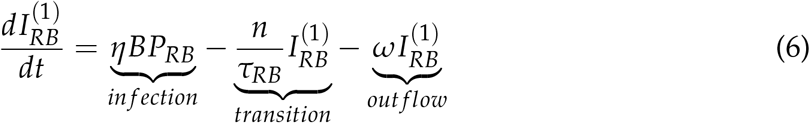

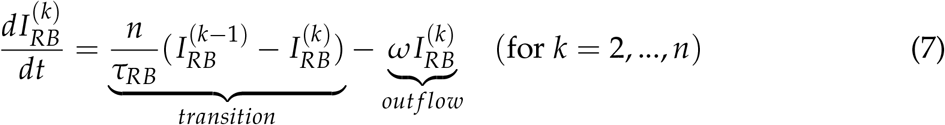

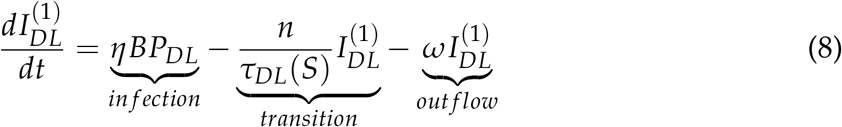

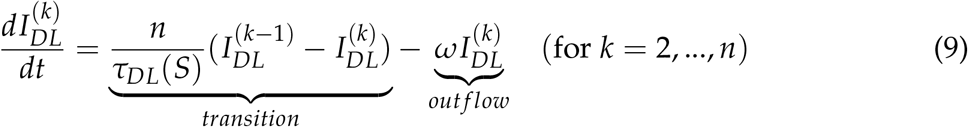

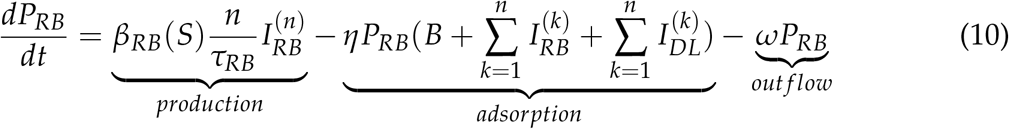

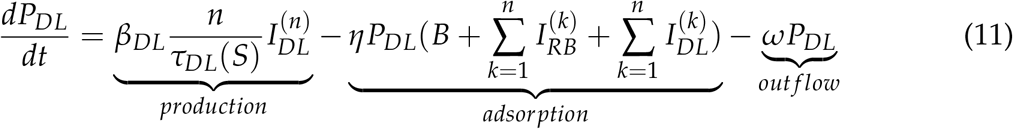

### Model 2 - Feast-Famine

To simulate a system with fluctuating substrate availability that alternates between phases of exponential growth and a stationary starvation phase, we modify the model as follows: We consider a batch culture system, where *B*_0_ bacteria, *P*_0_ phages, and *S*_0_ nutrients added to the system at *t* = 0. We simulate this system for one growth cycle that lasts *t*_*end*_ time units, which is set long enough in comparison to the initial bacteria and nutrient concentration that nutrients can be fully consumed and bacteria experience starvation (famine). At the end of one growth cycle, all bacteria and phages are diluted by a factor *δ*, which is typically set to *δ* = 1/10. Then, to restart a new “feast” phase, the concentration of the growth substrate is reset to the value *S*_0_ and a set amount of susceptible host bacteria *B*_*add*_ is added to the system. We repeat this procedure (usually 80 times) and follow the dynamics of the two phage populations.

During each cycle, the dynamics of the system are described by the same set of equations as the chemostat system, but with *ω* = 0. At the end of each cycle, the system components are adjusted in the following manner to serve as initial conditions for the next feast-famine cycle:

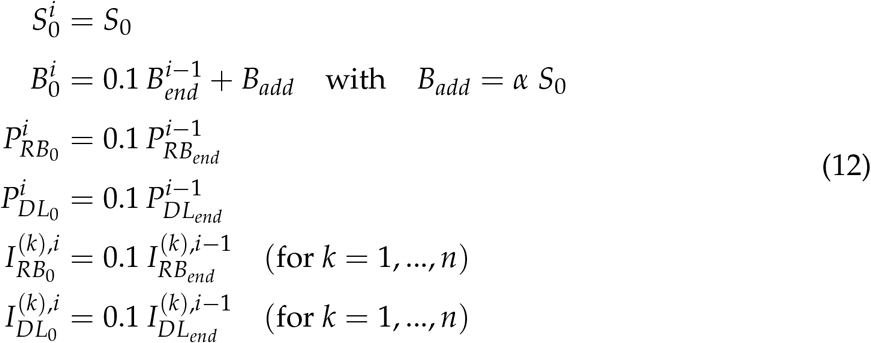

Here, the superscript *i* denotes the number of the feast-famine cycles, and the subscript 0 (*end*) denotes the start (end) of the next (previous) feast-famine cycles, respectively.

The Monod constant *K*_*s*_ in eq. (1) was set relatively low 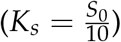 to permit rapid substrate consumption and entering of a starvation phase for a host population under attack. This simulates a true “feast” influx of excess nutrient which is rapidly consumed.

The numerical integration was performed using solve_ivp from the scipy.integrate package with the fourth-order Runge-Kutta method. A time unit *t* = *T*_*g*_ equals the generation time of the host bacterium at the maximum growth rate *µ*_*max*_, i.e., *T*_*g*_ = 1/*µ*_*max*_. For *E. coli* in rich medium, this corresponds to about 30 min. The maximum time step for the solver was set to 0.1.

**TABLE 1:** Default parameter values.

| Parameter | Description | Default value | Notes |
| --- | --- | --- | --- |
| $\mu_{max}$ | maximum bacterial growth rate | 1 | Time unit, see caption. |
| $K_s$ | half maximum growth constant | $10^5 \text{ ml}^{-1}$ | $K_s = S_0/10$ |
| $\eta$ | phage adsorption rate | $1.5 \cdot 10^{-8} \frac{\text{ml}}{T_g}$ | Adapted from adsorption rate reported for T4 [53]<br>$3 \cdot 10^{-8} \frac{\text{ml}}{h}$ |
| $\beta_{RB_{max}}$ | maximum burst size, phage RB | 150 | |
| $\beta_{DL}$ | burst size, phage DL | 150 | Adapted from burst size reported for T4 [53] |
| $\tau_{RB}$ | latent period, phage RB | $0.8 T_g$ | approximately 24 min, adapted from latent period reported for T4 [53] |
| $\tau_{DL_0}$ | adjusted minimum latent period, phage DL | $0.8 T_g$ | see $\tau_{RB}$ |
| $\gamma_{delay}$ | latent period delay responsiveness | 1.0 | |
| $S_{influx}$ | | $10^6 \text{ ml}^{-1}$ | only chemostat |
| $t_{end}$ | length of feast-famine cycle | 100 | only feast-famine |
| n | number of infection states | 5 | based on [54] |

**TABLE 2:**
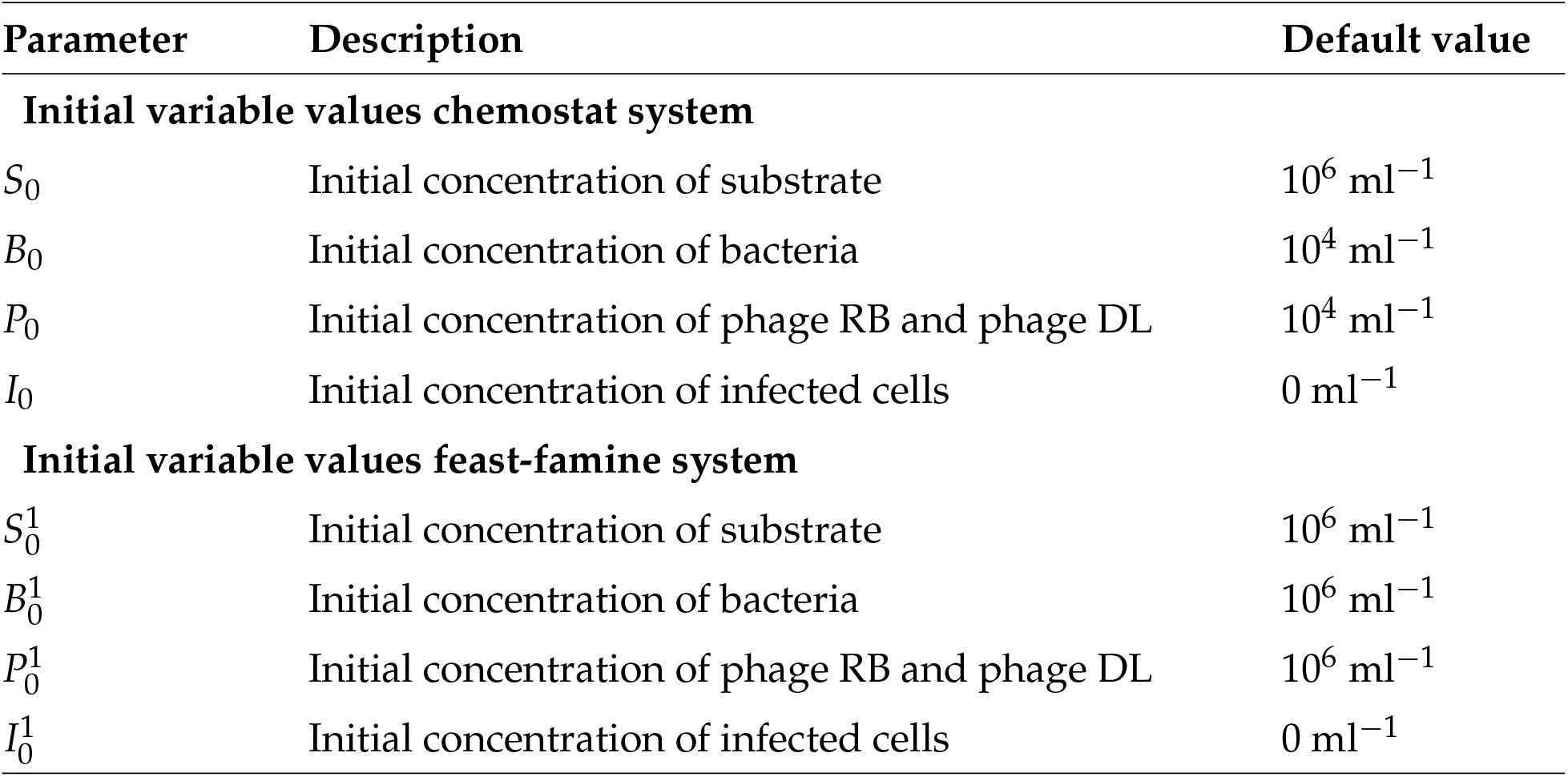
Initial values for the simulation.

## RESULTS

### In a chemostat system, delayed lysis gives only a limited competitive advantage

We first compete the two phages against each other in a chemostat system defined by equations (4)-(11). Important assumptions in the model are (i) phage RB has a reduced burst size if the substrate *S* < *S*_*th*_ = 2.5 · 10^4^ ml^−1^, (ii) phage DL has a prolonged latent period proportional to the host growth rate for the given *S*, and (iii) phages are adsorbed by both uninfected and infected cells, but the fate of the infection is determined by the primary infecting phage. Since phage RB’s latent period stays constant, the value of *γ*_*burst*_ up until which phage DL can outcompete phage RB gives a relative measure of the competitive strength of phage DL’s delayed lysis strategy in a given condition.

Figure **2**A shows one example of the population dynamics with the flow rate *ω* = 0.05 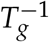 and a stationary phase burst size reduction factor for phage RB of *γ*_*burst*_ = 0.5. In the default case, we add the same amount of phage DL and phage RB at the start of the simulation (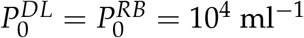). We observe oscillation of the system due to the negative feedback between the bacteria and phages, known as predator-prey oscillations. For this parameter set, the substrate concentration remains generally above *S*_*th*_, hence the burst size of phage RB is not reduced (eq. 10), while the latent period of phage DL increases periodically following the fluctuations in substrate concentration. This slight but continuous disadvantage leads to phage DL eventually being outcompeted by phage RB. The decrease in phage titer when the concentration of bacteria is low is mainly driven by the system flow rate.

**FIG 2:**
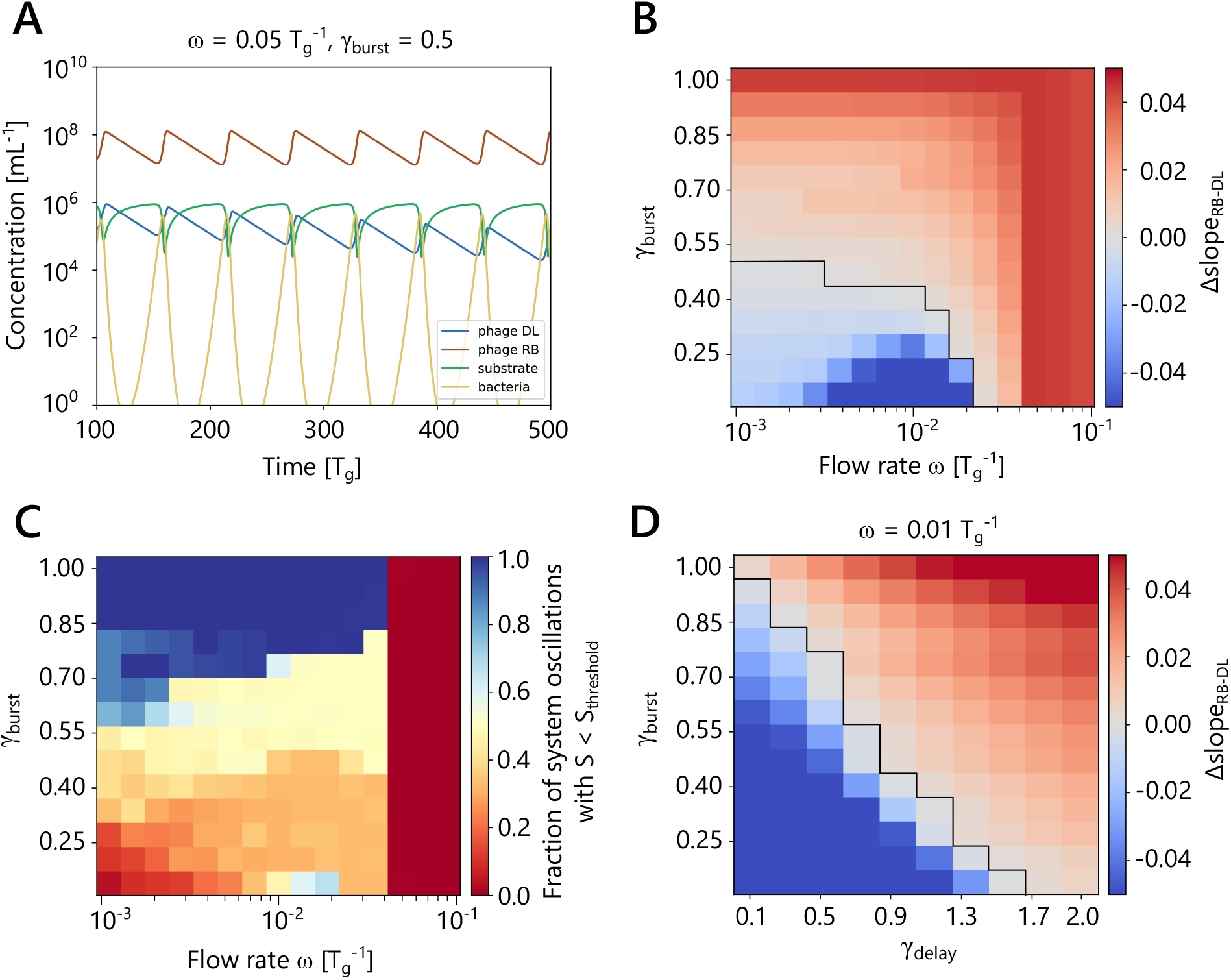
Simulation of lytic phage competition in a chemostat system A: Example of system dynamics, with flow rate *ω* = 0.05 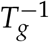 and phage RB burst size reduction factor *γ*_*burst*_ = 0.5. B: Outcome of phage competition in a chemostat environment for different values of *ω* and *γ*_*burst*_. For calculation of Δ*slope*, each simulation was evaluated for 50, 000 *T*_*g*_. A positive value (red) indicates that phage RB is outcompeting phage DL and *vice versa*. The black line highlights the border between positive and negative Δ*slope* outcomes. C: Relative number of oscillations in which the condition *S* < *S*_*th*_ = 2.5 · 10^4^ ml^−1^ is met, evaluated during the first 1/3 of the time series, normalized to total number of oscillations in that time period for each parameter set. D: Phase diagram screening the parameters *γ*_*delay*_ and *γ*_*burst*_, showing a clear tradeoff between the strength of the lysis delay response and burst size reduction. The black line highlights the border between positive and negative Δ*slope* outcomes.

In our system, the conditions for a reduced burst size of phage RB are only met for significant time periods when the flow rate *ω* is less than 0.04 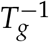 (Figure **2**C). Then phage DL can outcompete phage RB with *γ*_*burst*_ of up to 0.5, meaning that the burst size is reduced to 50% (Figure **2**B). Here, the outcome is defined as the difference of the slopes (Δ*slope*) of a log-linear fit to each phage concentration over time, normalized by the flow rate *ω*.

In the chemostat system, we can observe a direct tradeoff between increased latent period and reduced burst size under resource limitation. If we tune the intensity of the lysis delay response with the parameter *γ*_*delay*_ (eq. 2), while keeping the flow rate constant, we see that a weaker lysis delay response increases the competitive strength of phage DL relative to phage RB (Figure **2**D).

The boundary of the maximum *γ*_*burst*_ that phage DL can compete with at *ω* < 0.04 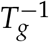 weakly depends on the initial conditions of the simulation. When we increase the initial concentration of one of the phages, the speed at which phage DL excludes phage RB in advantageous conditions is altered (Supplementary Figure S1). If phage RB is invading a system already dominated by phage DL, a value of *γ*_*burst*_ > 0.6 is necessary for successful invasion. For high flow rates (*ω* > 0.04 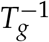), the nutrient *S* does not drop below the starvation threshold irrespective of the initial conditions and the outcome is unaffected. With all initial conditions tested, the effective latent period of phage DL is typically increased by 1.5 to 10 fold, and up to 20 fold in extreme cases at very low flow rates (Supplementary Figure S2). Overall, in the chemostat setup, the outcome of the competition is strongly limited by the flow rate and is not suitable to simulate prolonged periods of host starvation.

We modified the chemostat system by feeding bacterial host in addition to the growth substrate in different ratios (Supplementary Figure S3). This results in a dampening of the oscillation of the system and removes environmental fluctuation, creating a more stable environment, even with a very low bacterial concentration in the influx. Except for certain edge cases, the system reaches a steady state with both phages present, with *P*_*RB*_ > *P*_*DL*_.

### Repeated feast-famine cycles increase the fitness of a delayed lysis strategy

In most natural ecosystems, substrate influx is not continuous and bacteria spend much of their time in nutrient depleted conditions [15, 16, 17, 18, 19]. We therefore next compare the infection dynamics of the two phages targeting a shared host population in a series of feast-famine cycles. We consider a batch culture where a limited amount of nutrients supports populations of bacteria and phages. After a set time period, the end product of this cultivation (phages, bacteria (uninfected and infected), and the remaining substrate) is diluted using a fixed dilution factor (1/10) and added to a new batch culture. A growth cycle lasts for 100 *T*_*g*_ time units, equivalent to approximately 48 hours for a fast growing bacterium like *Escherichia coli*. To restart a new “feast” phase, the concentration of the growth substrate is reset to *S*_0_ and a set amount of susceptible host bacteria *B*_*add*_ is added to the system (Figure **3**A). We define the ratio of *B*_*add*_ and *S*_0_ as *α* (eq. (12)), and explore the effect of *α* on the competition. Since the nutrient is measured as a unit required for doubling of one bacterium, 1/*α* = *S*_0_/*B*_*add*_ quantifies the potential bacterial growth fold per feast phase and therefore also sets the timeline until the starvation phase is reached, with larger *α* values leading to shorter growth phases (Supplementary Figure S4A). Typically, we simulate 80 such feast-famine cycles for each parameter combination.

**FIG 3:**
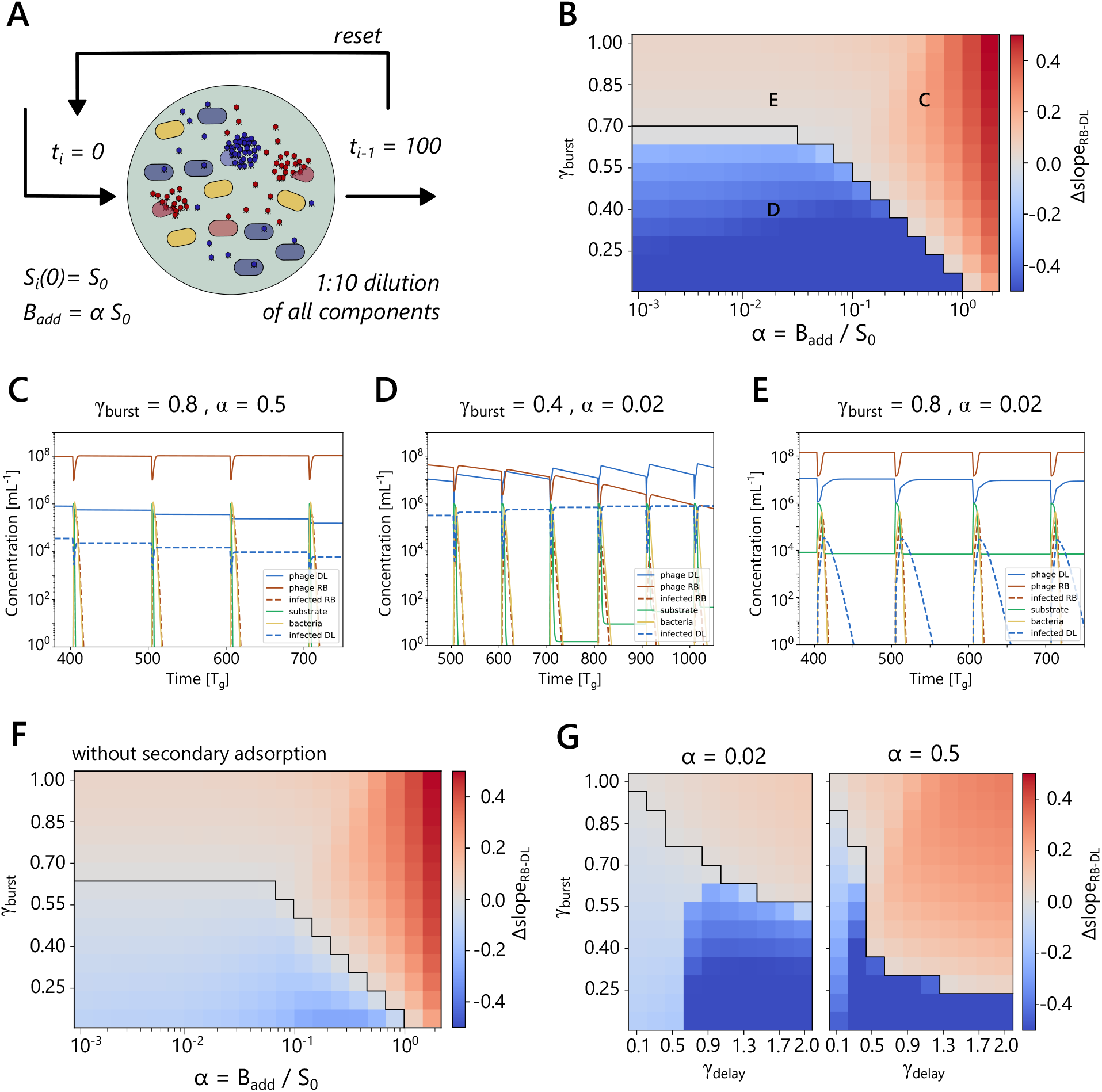
Simulation of phage competition in a series of batch cultures (feast-famine system) A: Schematics illustrating the feast-famine system. B: Result of phage competition in the feast-famine system for different values of *γ*_*burst*_ and *α*. The result is shown as the difference of the slopes of a log-linear fit to each phage concentration over time. A positive value (red) indicates that phage RB is outcompeting phage DL and *vice versa*. See details in Supplementary Figure S6. The black line highlights the border between positive and negative Δ*slope* outcomes. Letters indicate the parameters used for the examples shown in panel C-E. C: Example dynamics of the feast-famine system (cycle 4 - cycle 7) for *γ*_*burst*_ = 0.8, *α* = 0.5. D: System dynamics of the feast-famine system (cycle 5 - cycle 9) for *γ*_*burst*_ = 0.4, *α* = 0.02. E: System dynamics for *γ*_*burst*_ = 0.8, *α* = 0.02 (cycle 4 - cycle 7). F: Result of phage competition in the feast-famine system for different values of *γ*_*burst*_ and *α* as in B, but without the term for phage adsorption to infected cells in eqs. 10 and 11. The black line highlights the border between positive and negative Δ*slope* outcomes. G: Sensitivity of the competition outcome on the strength of the lysis delay response parameter *γ*_*delay*_ for *α* = 0.02 and *α* = 0.5. The black line highlights the border between positive and negative Δ*slope* outcomes.

We screened the dynamics of the feast-famine system for different values of the parameters *γ*_*burst*_ and *α* and observed a number of possible outcomes (Figure **3**B). We characterize the outcome by the difference between slopes of a long-term population log-linear fit; negative values (blue) denote that phage DL outcompetes phage RB, and *vice versa* for positive values (red). The details of the log-linear fit and the calculation of Δ*slope* are given in Supplementary Figure S6.

The tested parameter space can be separated into two main regions that produce the outcome of the competition by different mechanisms. The border between these two main regions lies approximately at *α* = 0.1. For *α* < 0.1 the competitive strength of phage DL only weakly depends on *α*, while for *α* > 0.1, we observe a clear dependence of the competitive strength of phage DL on *α*. High *α* values (*α* > 0.1) lead to a rapid onset of the starvation conditions. Without phages present, the time to consume all of the substrate *S*_0_ by newly added bacteria *B*_*add*_ = *αS*_0_ would be *T*_*sub*_ = ln(1 + *α*)/*αT*_*g*_, which satisfies *T*_*sub*_ < 2.5 *T*_*g*_ for *α* > 0.1. Even though the presence of phages somewhat slows down substrate consumption, the extremely short growth window in the feast period results in a disadvantage for phage DL. Under these conditions, the lysis of the carried-over population of phage DL-infected cells cannot proceed efficiently, and any newly infected cells are immediately affected by a strong lysis delay (see example dynamics for *γ*_*burst*_ = 0.8, *α* = 0.5, Figure **3**C). Hence, phage DL loses the opportunity to build a large population of free phage to compete against phage RB in claiming susceptible hosts, and this disadvantage becomes more severe the shorter the growth period is. Phage RB is also affected by the short growth window, leading to many RB-infected cells bursting with the reduced burst size *γ*_*burst*_ 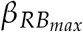. Larger *α* values lead to shorter growth windows, which means that phage DL gets outcompeted by phage RB with progressively smaller *γ*_*burst*_ values. This results in a downward slope of the boundary between phage DL and phage RB advantageous parameter regions in the *α*-*γ*_*burst*_ space (Figure **3**B).

When *α* < 0.1, it takes longer for newly added bacteria to consume all the substrate, opening up for more complex dynamics. This is exemplified in Fig. **3**D for *γ*_*burst*_ = 0.4 and *α* = 0.02. The substrate decreases as the bacteria grow after the system reset. Then the concentration of susceptible bacteria drops rapidly due to phage infection. Those infected by phage RB lyse quickly and produce new phage RB, while the concentration of bacteria infected by phage DL plateaus as the substrate concentration decreases towards its minimum level typically in less than 7 *T*_*g*_ time units, and lysis comes to a halt. In these conditions the lysis will be delayed about 10 times the length of the feast-famine cycle period, effectively stopping any lysis until the system resets. The decrease in both phage populations until the next dilution event is mainly driven by adsorption to DL-infected cells that are still in the system. At the end of the cycle, phage RB, phage DL, and DL-infected bacteria are present in the system, and 1/10 of them continue to the next feast round (eq. (12)). After the system reset, the DL-infected cells experience nutrient rich conditions and lyse quickly while the susceptible host *B* is at its maximum concentration. The freshly produced phages can capture a large amount of the newly available host population and thus slowly outcompete phage RB.

However, when *α* < 0.1 and *γ*_*burst*_ > 0.7, phage DL is outcompeted by phage RB irrespective of *α*. This happens because, a high value of *γ*_*burst*_ increases the concentration of phage RB after the first cycle (Supplementary Figure S5A). The elevated phage titer leads to increased killing of the relatively small host population and therefore higher residual substrate concentrations. In these conditions, no substantial latent period increase is initiated, preventing the DL-infected cells from carrying over to the next growth cycle and releasing new phages early instead (Figure **3**E). We analyzed the typical latent period experienced by DL-infected cells in this parameter region and found that for *γ*_*burst*_ > 0.7 and *α* < 0.1 DL-infected cells on average experience a less than 32 *T*_*g*_ latent period, meaning that all infected cells will progressively lyse before the system reset happening every 100*T*_*g*_ (Supplementary Figure S4B).

It is worth noting that the lysis delay gives secondary advantage to phage DL. Adsorption to DL-infected unlysed cells act as an effective sink for free phages, inactivating both phage RB and phage DL particles. If we remove the term for secondary, abortive adsorptions from the model, we see a strong impact on the ability of phage DL to rapidly exclude phage RB from the system in DL-favorable conditions (Figure **3**F). While the overall boundary of favorable conditions for each phage shifts only marginally, the rate at which phage DL can competitively exclude phage RB from the system is heavily diminished, especially for *α* < 0.1. We can further analyze the importance of a sufficiently strong lysis delay for the competitiveness of phage DL by modulating the *γ*_*delay*_ parameter. For *α* < 0.1 (exemplified by *α* = 0.02 in Figure **3**G) there is a sharp transition to the ability to rapidly exclude phage RB with *γ*_*burst*_ ≤ 0.7 at around *γ*_*delay*_ = 0.6. This dependence does not exist for *α* > 0.1. This leads us to conclude that two dynamics linked to the lysis delay are key in supporting the competitiveness of phage DL in a feast-famine system, specifically when *α* is low: a sufficiently strong lysis delay enables reduction of free phage during the starvation phase through secondary, abortive adsorptions and enables infected cells to wait for optimal conditions to lyse at a later time. Note that phage loss by adsorption is not as important in the chemostat system in the studied parameter range, instead the phage decay is mainly driven by the continuous flow (Supplementary Figure S2B).

The position of the boundary between phage RB and phage DL exclusion depends on the initial conditions of the system. In Figure **3**B, We used the same initial conditions in all cases: *P*_0_ = *B*_0_ = *S*_0_ = 10^6^ ml^−1^, effectively setting *α* = 1.0 for the first cycle. Due to the immediate consumption of the substrate by the initial host population, most of the RB-infected cells will lyse with the reduced burst size, leading to a *γ*_*burst*_ dependent phage RB concentration at the end of cycle 1 (Supplementary Figure S5A), which then determines the dynamics in the following cycles. If we instead initialize with an increased concentration of phage DL (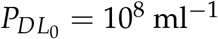), the bacterial growth and substrate consumption dynamics in the first few cycles are changed in a way that activates delayed lysis and carry-over of DL-infected cells even for large *γ*_*burst*_ values (Supplementary Figure S5C), extending the same dynamics that are observed for larger *α* values. In the reverse case of setting 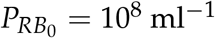, the outcome is largely the same as with the default initial conditions (Supplementary Figure S5B), highlighting that we are effectively analyzing the ability of phage DL to invade an established population of phage RB in our default setup.

### Fluctuating environments support the coexistence of competing phages

#### The effect of pairwise alternating host feeding rates

The feast-famine system introduces a cyclic variation between periods of rapid microbial growth and starvation. However, the variation follows a regular pattern that favors either one of the phages in the system and leads to gradual competitive exclusion of the other. We next introduce additional environmental variation in the system by either alternating between two *α* values (mode II in Figure **4**A) or by choosing a random *α* value at each reset event (mode III). We investigate how the different parameter regimes affect the fitness of each phage species and if it allows for long-term coexistence of phage DL and phage RB.

**FIG 4:**
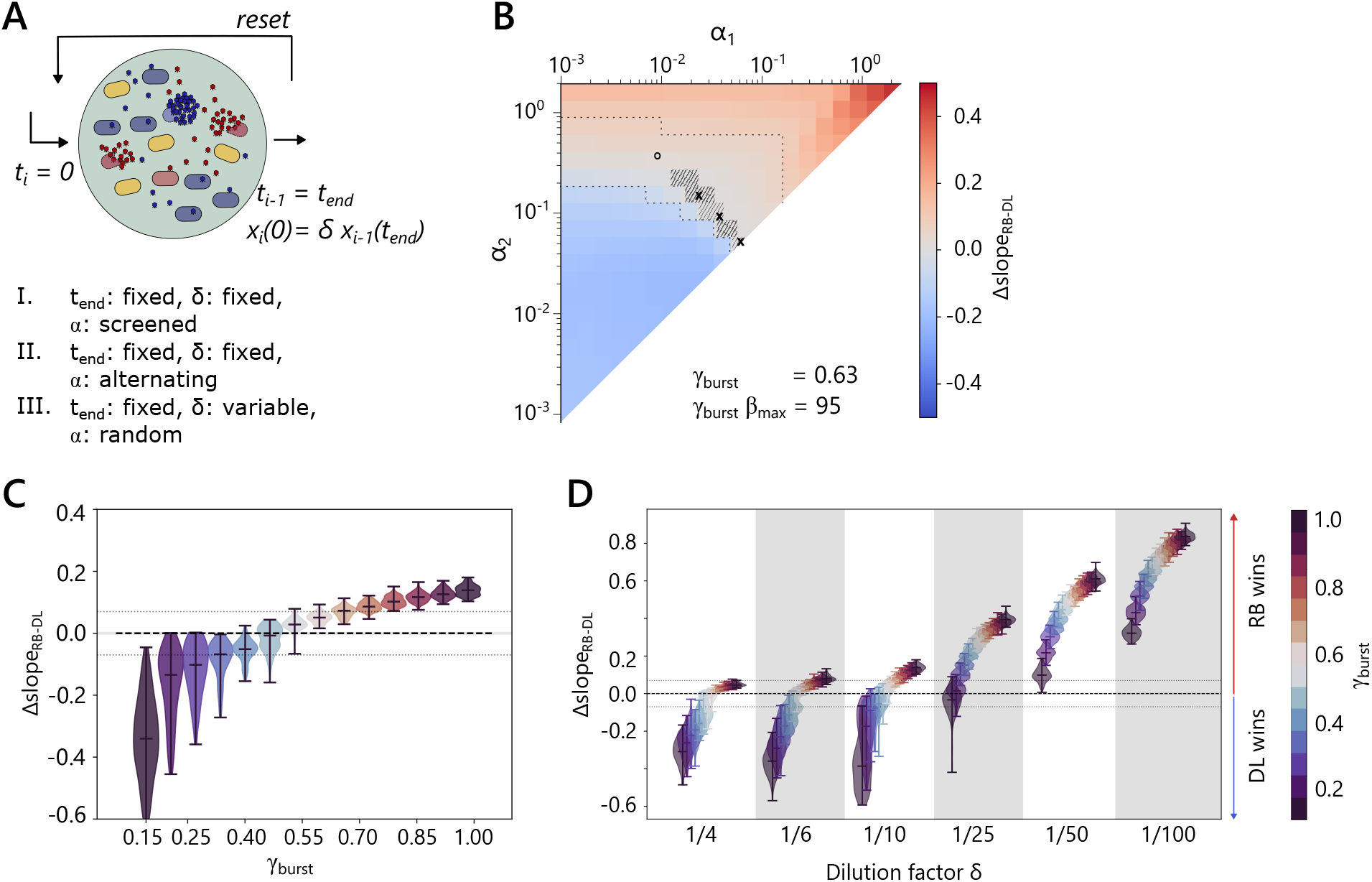
A: Overview of different modes of controlling the environment in the feast-famine system. B: Results from mode II. Alternating between a pair of *α* values broadens the parameter range in which the two phages can coexist. Cross: At *α* = 0.057, *γ*_*burst*_ = 0.63 the two phages coexist over the simulated time scales. Certain pairs of *α*_1_ and *α*_2_ lead to coexistence (shaded area, black crosses). In those cases the average *α* is still 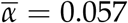. This finding is, however, not generalizable for the whole parameter space (open circle indicates counterexample). C: Results from mode III, drawing a random *α* value at each reset. Each violin plot represents the results from 100 simulations using a randomly selected value for *α* at each reset event and a specific stationary phase burst reduction factor for phage RB. Dotted lines indicate the thresholds set for defining slow exclusion of the competitor phage. D: The dilution factor at the reset event is an important environmental variable of the system. As in C, which shows a more detailed version of the simulations using *δ* = 1/10, each violin plot shows the results from 100 simulations using a series of randomly selected *α* values. Phage DL can only competitively exclude phage RB in systems with sufficiently low dilution factors, akin to lower flow rates in the chemostat system.

There is a vast foundation of both theoretical and experimental research that shows that environmental fluctuations can stabilize the coexistence of species which could otherwise not thrive alongside each other [55, 56, 57, 58]. The key mechanisms for the fluctuation-dependent coexistence include the presence of relative nonlinearities in the response to the fluctuating environment [59, 60, 61] and the storage effect [62]. Often, multiple mechanisms are contributing to coexistence simultaneously [63, 64, 65]. In some systems, the coexistence in a fluctuating environment may be explained by fluctuation-independent mechanisms, i.e., the coexistence in a fluctuating environment can be predicted by the coexistence in the time-averaged, constant environment [66, 67]. The relevance of each mechanism depends on the system at hand, and specifically if the per capita growth rate is dependent on competitive factors in a linear or nonlinear manner. Our current model assumes nonlinear substrate dependence of the phage growth rate through either burst size or latent period variation, and it is not trivial to map the model to previously studied analytically tractable models [66, 67]. Hence, we here study the model outcome in fluctuating environments numerically to highlight if and how phage coexistence can be realized in fluctuating environments.

First, we evaluate a regime of environmental fluctuation in which the parameter *α* alternates between two values *α*_1_ and *α*_2_ within the previously tested parameter range. Interesting dynamics can be found for *γ*_*burst*_ = 0.63 (Figure **4**B), since this parameter range captures a broad range of conditions where the competitive strength of phage DL and phage RB is approximately equal and there is potential for coexistence of the two phages in specific conditions. The diagonal cut-off line of this plot is equivalent to the corresponding horizontal section in Figure **3**B showing data for a non-alternating *α* value.

When *γ*_*burst*_ = 0.63 and *α* = 0.057, (cross symbol on the diagonal), the system outcome falls into the defined range of coexistence and both phages can coexist in the feast-famine system over the tested time scales (Supplementary Figures S7A and S6 for a definition of coexistence in this model). On this basis, we can then construct cases with alternating *α* values where the geometric mean falls on *α* = 0.057. Examples are (*α*_1_ = 0.09, *α*_2_ = 0.036) or (*α*_1_ = 0.143, *α*_2_ = 0.023), which are shown in Figure **4**B as cross symbols.

We use the geometric mean as the effective constant *α* because, in the Lotka–Volterra system studied by Abreu et al. [67], alternating dilution factors produce long-term dynamics equivalent to those under a constant dilution equal to the geometric mean. In our model, *α* plays an analogous multiplicative role, setting the fold-change permitted by the ratio of initial nutrients to initial bacterial density. Although Abreu’s analytical equivalence relies on the linear structure of the LV equations and does not directly extend to our more nonlinear system, the geometric mean remains the natural choice for comparing alternating and constant-parameter scenarios.

The shaded area in Figure **4**B indicates the broader range where we find pairs of *α* values that result in coexistence. We find that switching between two *α* values with the geometric mean in the range of 0.056 < *α* < 0.06 results in phage coexistence over the tested time scales (at different phage titers, see details in Supplementary Figure S7B, C). However, this effect is quite unstable. Pairs of *α* values further away from the target mean do not result in coexistence (e.g., *α*_1_ = 0.36, *α*_2_ = 0.009, open circle in Figure **4**B and Supplementary Figure S7D).

#### Randomly selected host feeding rates allow a broader examination of environmental parameters

We showed that alternating between two distinct conditions can promote coexistence of the two phage types. We next extend this idea by allowing *α* to vary at random (option III in Figure **4**A). At every reset event, a random value of *α* from within the range 0.001 ≤ *α* < 2 is uniformly drawn. We ran 100 simulations for each of the 14 values of *γ*_*burst*_. The results are presented as distributions of Δ*slope* outcomes in Figure **4**C. As before, a positive Δ*slope* value means that phage RB has an advantage over phage DL. Under these conditions, we see that phage DL can only outcompete phage RB if *γ*_*burst*_ < 0.5. We also found that phage RB and phage DL can frequently coexist over long time periods when 0.4 < *γ*_*burst*_ < 0.6 (Example dynamics shown in Supplementary Figure S8).

The random draw of a host feeding rate *α* removes this parameter as the key environmental variable, and leaves the dilution factor at each reset event as another controllable parameter, which has been kept constant as 1/10 so far. To evaluate the role of this parameter in determining the outcome of the phage competition, we systematically changed the dilution factor and simulated the competition outcome (Figure **4**D). The dilution factor corresponds to a system wide mortality rate that effects both phages and hosts equally. A high dilution factor, representative of a very harsh external environment, favors the faster replicating phage RB, just like a high flow rate *ω* is favorable for phage RB in the chemostat system. The length of the starvation period also impacts the outcome of the competition, since abortive adsorption of phages to already infected cells during this period is an important contribution to phage DL fitness (see Figure **3**). However, the effect of increasing or decreasing the time scale of each feast-famine cycle is much smaller than the effect of changing the dilution factor (Supplementary Figure S9A). In the very extreme case of frequent, small dilutions (*t*_*end*_ = 10, *δ* = 0.9) the feast-famine system approaches the case of the chemostat system with a host feeding rate, i.e. relatively stable coexistence due to reduced host and substrate fluctuation (Supplementary Figure S9B, compare with Supplementary Figure S3)

In the previous examples, we have seen that the outcome of the competition with alternating *α* values can in some cases be predicted by analyzing the system with the averaged, static *α* value, pointing towards fluctuation-independent mechanisms determining the system outcome [67]. This observation was, however, only made within a limited parameter range. In the case of the random selection of *α* values, we observe clear deviation from the outcome of simulations with a static, averaged *α* value (Supplementary Figure S10).

## DISCUSSION

Just like any other living organism, bacteriophages must evolve strategies to overcome adverse environmental conditions [22] and compete with other organisms that occupy the same ecological niche. Phage evolution and growth strategies are often evaluated under the premise of co-evolution in an arms race with their host [8, 3, 9, 10]. However, before they encounter potential host defenses, they first have to compete with other phages in claiming a susceptible host.

In this study we explore the dynamics between two virulent phages infecting the same host population under the pressure of varying availability of the bacterial growth substrate. We find that repeated feast-famine cycles improve the competitive strength of a phage that has a mechanism to delay lysis in response to host starvation against a rapidly lysing phage with a reduced burst size, and we identified several mechanisms that contribute to this phenomenon. We also find that increased variation in the environmental conditions can support the long term co-existence of two phage types with different lysis behaviors.

The direct implementation of feast-famine cycles was necessary to observe these effects. In a continuous system like the chemostat system analyzed in part 1, fluctuations are generated due to the predator-prey dynamics of the bacteria-bacteriophage mixture. In the tested parameter range we then observe relatively stable oscillations with both phages present over long periods over time, but usually one of them is trending towards extinction from the system eventually. These fluctuations, however, do not generate lasting starvation periods, since the growth substrate is continuously replenished. Similarly, the addition of susceptible host at the start of a new feast-famine cycle is necessary to observe certain specific advantages of the lysis delay strategy, such as rapid lysis of previously infected cells when a new pool of nutrients and hosts becomes available. We can easily imagine natural scenarios with such dynamics, e.g. availability of a previously spatially separated subpopulation and nutrients after a washout/rainfall event or introduction of both nutrients and bacterial hosts after a meal in the animal gut.

Our focus point for this study is to understand under which environmental conditions lysis delay strategies produce the biggest fitness effect for a bacteriophage. In order to measure the competitive strength of phage DL, we introduce a competitor phage RB, which instead of delaying lysis, will suffer a reduced burst size when the host physiology is sufficiently diminished.

We assume here that *τ*_0_ and *β*_*max*_ constitute the optimal parameter set for these phage types at the maximum growth rate. Phage DL then reacts very sensitively to changes in the environment and in the host growth rate by delaying lysis in order to still yield the maximum burst size in a slower growing cell. We assume that the control phage RB has the ability to maintain the optimal lysis behavior in a broader range of environmental conditions, and is only forced to reduce the burst size when the host growth as slowed down significantly. These relationships do not directly follow experimentally tested data, but are loosely inspired by the behavior of two well-studied model phages of *Escherichia coli* (T4 and T7). The variety of system dynamics we observe even with this simple model demonstrates the need for more experimental study of phage response to host starvation and determination of phage replication parameters along the whole growth curve. Many of these experiments would be very laborious with the current standard methodologies, especially if systematic study across many conditions is planned. This prompts the development of new high-throughput methodologies for phage characterization (cf. [54]), which are trailing behind compared to methods for characterization of bacteria [68].

Despite the limitations, the overall qualitative outcomes of this study are experimentally testable. We can think of setting up a series of batch cultures equivalent to the feast-famine system presented. If such a system were simultaneously infected with a phage with a delayed lysis mechanism (e.g. T4) and a continuously lysing phage (e.g. T7), the winner of the competition following different feeding schemes could be determined by classical plaque assays, due to the difference in plaque morphology of these two phages. It should, however, be noted that the experimental testing would be complicated by the development of resistance to phage infection, which was not considered in the model. Development of resistance and therefore divergence in the host population and phage host range can increase the potential for coexistence of similar phages [69, 70, 71, 7]. In the model, we can tune the fitness of the competitor phage to test the limits of the fitness of the delayed lysis phage and its response to various environmental disruptions in a very fine-grained matter. An experimental test will be limited by the replication characteristics of available model phages.

Our findings on the possibility for long term coexistence of competitor phages in fluctuating environments evoke some similarity to findings of classical ecological modeling concerning coexistence of species on shared resources. The delayed lysis/pseudolysogeny could be interpreted as a form of buffered population growth, one of the essential contributors to coexistence through the storage effect [62]. Our model does not directly fit into the theoretical frame work of analytically trackable modeling concerning coexistence on shared resources [66, 67], e.g. we assume that interspecies and intraspecies competition for the limited resource has the same magnitude, but we observe some of the hallmarks of both fluctuation-dependent and fluctuation-independent mechanisms driving coexistence in the conditions where we observe it. We found examples where the geometric mean of an alternating pair of *α* values could predict the outcome of the competition, even though explicit nutrient dependence makes the current model highly nonlinear. This observation is consistent with the previously reported usefulness of such approach in understanding complex communities [67] and can be interpreted as a good approximation if the fluctuation amplitude is sufficiently small.

The delayed lysis mechanism of phage T4, which served as an inspiration for phage DL in the model, is a very intricate system that gives phage T4 multiple options to regulate its response to environmental and host information. The classically known mechanism is lysis inhibition, where lysis halts in response to adsorption of multiple of the same phage [31]. Delayed lysis was also observed after T4 infection of non-growing, starved cells, independent of MOI [30, 72]. In our model, we did not consider secondary-infection triggered lysis inhibition, but we observed that the adsorption of free phage particles to already infected cells is crucial for the fitness of a delayed lysis phage in a feast-famine environment. Interestingly, in our previous theoretical work of competition between a lysis inhibiting wild type T4 phage and a non-lysis inhibition mutant phage, we also reported that adsorption of phages to the lysis inhibited cells contribute to the competitiveness of the wild type T4 phage [73]. We therefore interpret the lysis delay strategy not just a mechanism for the phage to hibernate in an environment when the outside conditions might be harsh for free phage particles, but as an active competitive ability to modify the environment by reducing both interspecies and intraspecies competition in poor resource conditions. Such a mechanism is of course beneficial only if the secondary adsorption does not prevent the eventual production of primary infecting phage. We therefore hypothesize that the ability to delay lysis for a virulent phage should in many cases co-occur with mechanisms of superinfection exclusion across different phage species.

The ability to modulate lysis timing in response to host physiology comes with a cost to short-term replication but can substantially enhance phage persistence in resource-limited and fluctuating environments. Our simulations highlight the specific ecological conditions under which delayed lysis is favored or suppressed, and show how the presence of competing phages reshapes infection and growth dynamics for the entire system. These findings underscore that phage life-history traits cannot be evaluated in isolation from the surrounding biotic and abiotic context. A deeper understanding of how physiological cues, competition, and environmental variability jointly govern phage replication strategies will be essential for improving predictive models of microbial ecosystems and for guiding the rational design of phage cocktails in therapeutic settings.

It is known that bacterial growth dynamics are modulated by the presence of phages. We show here that this modulation does not require specific interactions with the host metabolism, but that changes in the composition of the phage population and small differences in the infection dynamics of the phages in the community impact substrate consumption dynamics and bacterial population growth. Implementing these results in a broader context of studying microbiome dynamics underscores that phages are a key component of ecosystems and influence nutrient fluxes, community composition, and growth dynamics.

## Supporting information

EhrmannMitarai_2026_SupplementaryFigures

## DATA AVAILABILITY STATEMENT

All code used for simulations and figure generation is available on GitHub https://github.com/AnjaEh/Ehrmann_Mitarai_2025_FeastFamine

