## Supplementary material for "Host Physiology–Dependent Lysis Timing Shapes Bacteriophage Competition Under Nutrient Fluctuations": EhrmannMitarai_2026_SupplementaryFigures

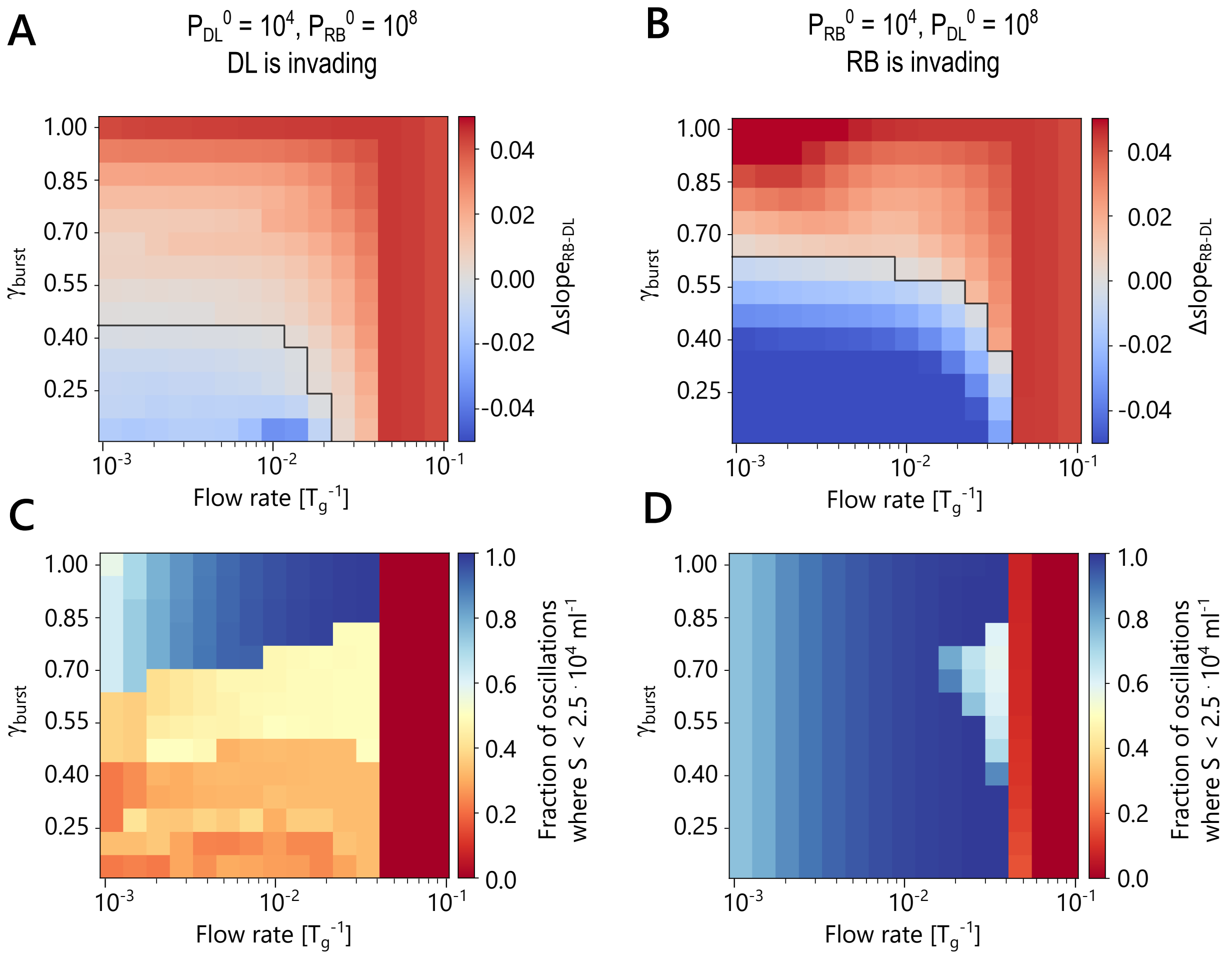


**SUPPLEMENTARY FIGURE** S1 Outcome for the chemostat system with different initial conditions. A: Phage DL is invading a system with dominant phage RB. Where phage DL has an advantage, the rate at which it can exclude phage RB is slower, but otherwise the outcome is the same as with the default initial conditions. B: phage RB is invading a system with dominant phage DL. When phage DL starts as a dominant species, it can outcompete phage RB with a higher $\gamma_{burst}$ value, under the condition that the flow rate $\omega < 0.04$. For $\omega > 0.04$ the substrate concentration does not drop below the threshold $S(t) < S_{th} = 2.5 \cdot 10^4$ ml$^{-1}$ under these conditions (panel D). In both A and B, $\Delta slope$ is normalized by the flow rate. C, D: fraction of system oscillations in which the substrate concentration drops below the starvation threshold $S(t) < S_{th} = 2.5 \cdot 10^4$ ml$^{-1}$, based on the first third of the system oscillations in the time series for each of the system invasion conditions, respectively. E, F: for each of the system invasion conditions, respectively.

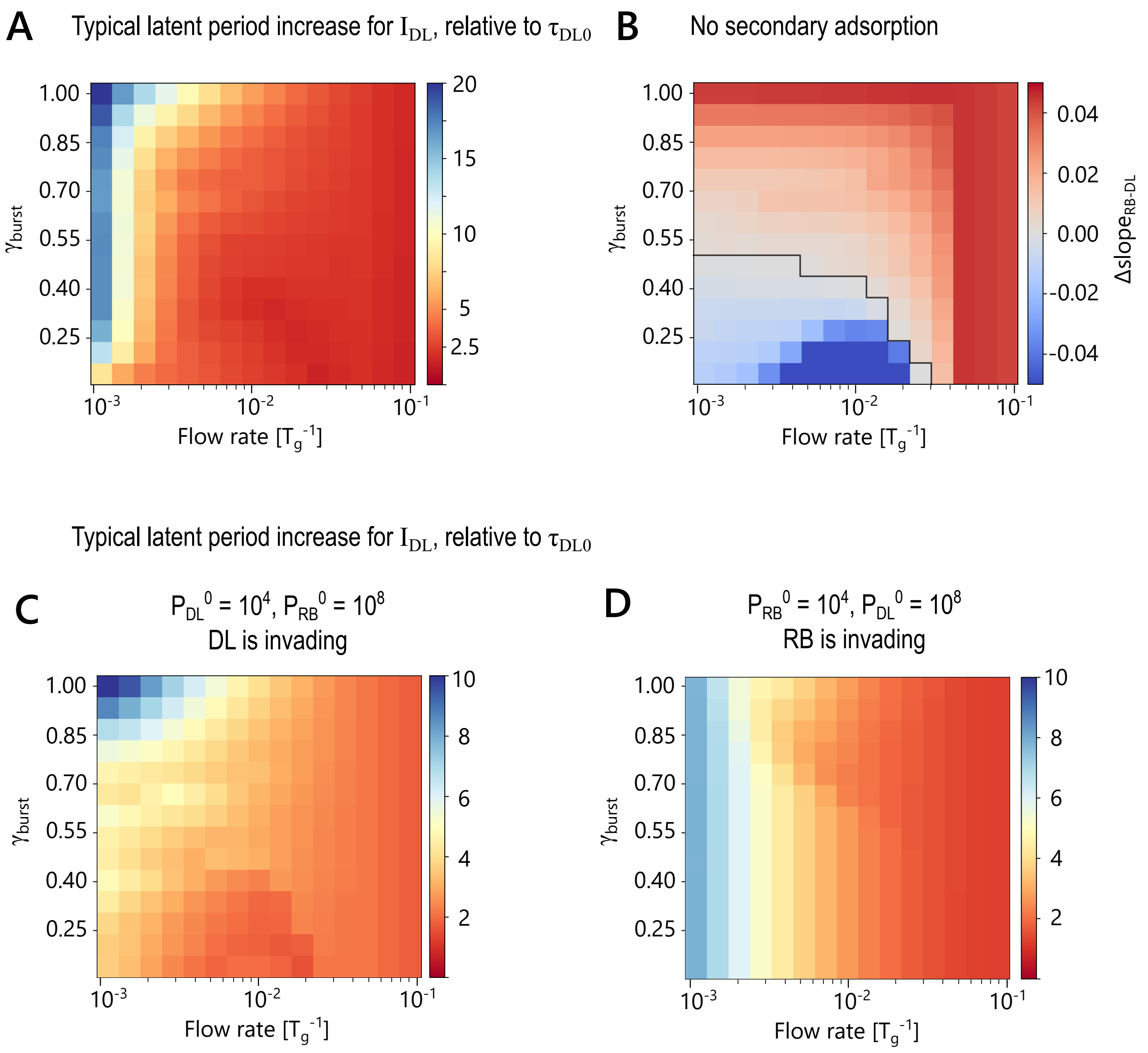


**SUPPLEMENTARY FIGURE** S2 Effective latent period increase of phage DL infected cells and importance of secondary adsorptions in the chemostat system. A: The typical latent period increase relative to $\tau_{DL_0}$ experienced by $I_{DL}$. Calculated as the weighted mean of $\tau_{DL}(S)$ over the time intervals where $\sum_{k=1}^{n} I_{DL}^{(k)} > 1$. B: Outcome of the chemostat system if the terms for adsorption to already infected cells are removed from eqs. 11 and 10. C, D: The typical latent period increase relative to $\tau_{DL_0}$ experienced by $I_{DL}$, as described for panel A, but with adjusted initial conditions to simulate system invasion by either phage. Compare with S1 and note that the scaling of the colormap is different in panel A.

with alternative eq. (5)

$$\frac{dB}{dt} = \underbrace{\omega(B_{influx} - B)}_{in/outflow} + \underbrace{\mu_{max}\frac{S}{K_s + S}B}_{cell\ division} - \underbrace{\eta B(P_{RB} + P_{DL})}_{infection} \quad , \qquad \alpha = B_{influx} / S_{influx}$$

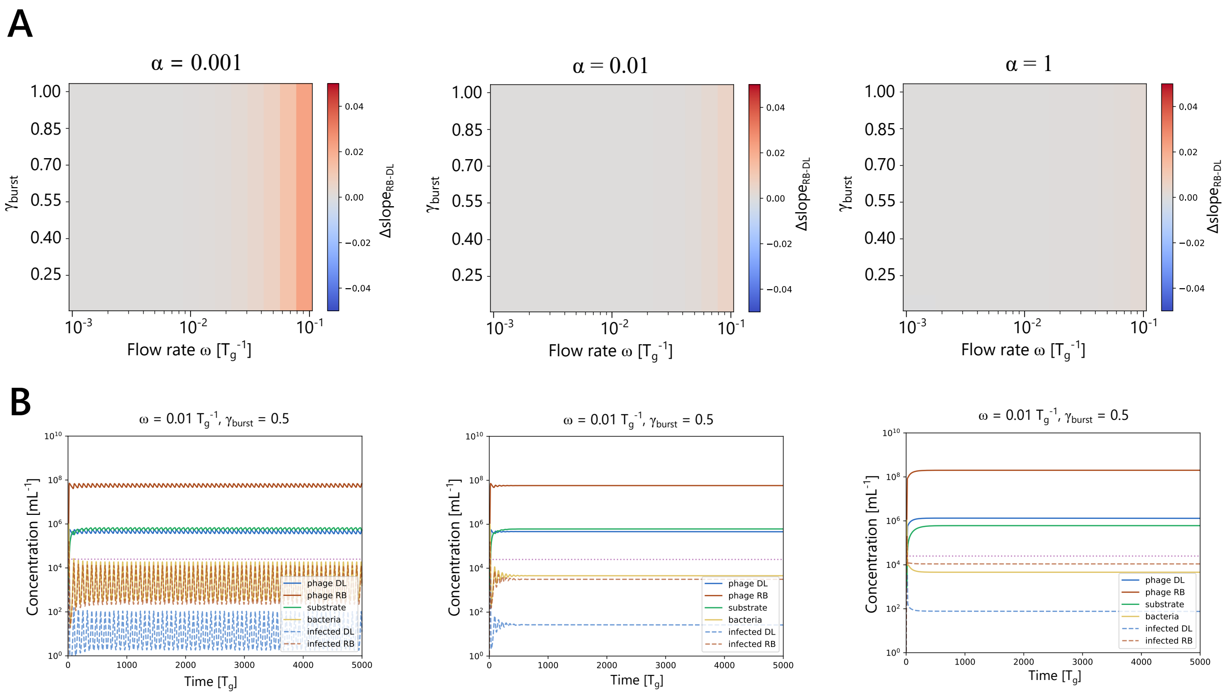


**SUPPLEMENTARY FIGURE** S3 Chemostat system with additional influx term of susceptible host bacteria. The system is modified to include an additional influx term of susceptible host bacteria in eq. 5. A: Outcome of the competition between phage DL and phage RB in the modified system for different values of $\alpha = B_{influx}/S_{influx}$. $\Delta slope$ is normalized by the flow rate $\omega$. B: Example dynamics matching each of the host feeding rates in panel A at $\omega = 0.01 T_g^{-1}, \gamma_{burst} = 0.5$

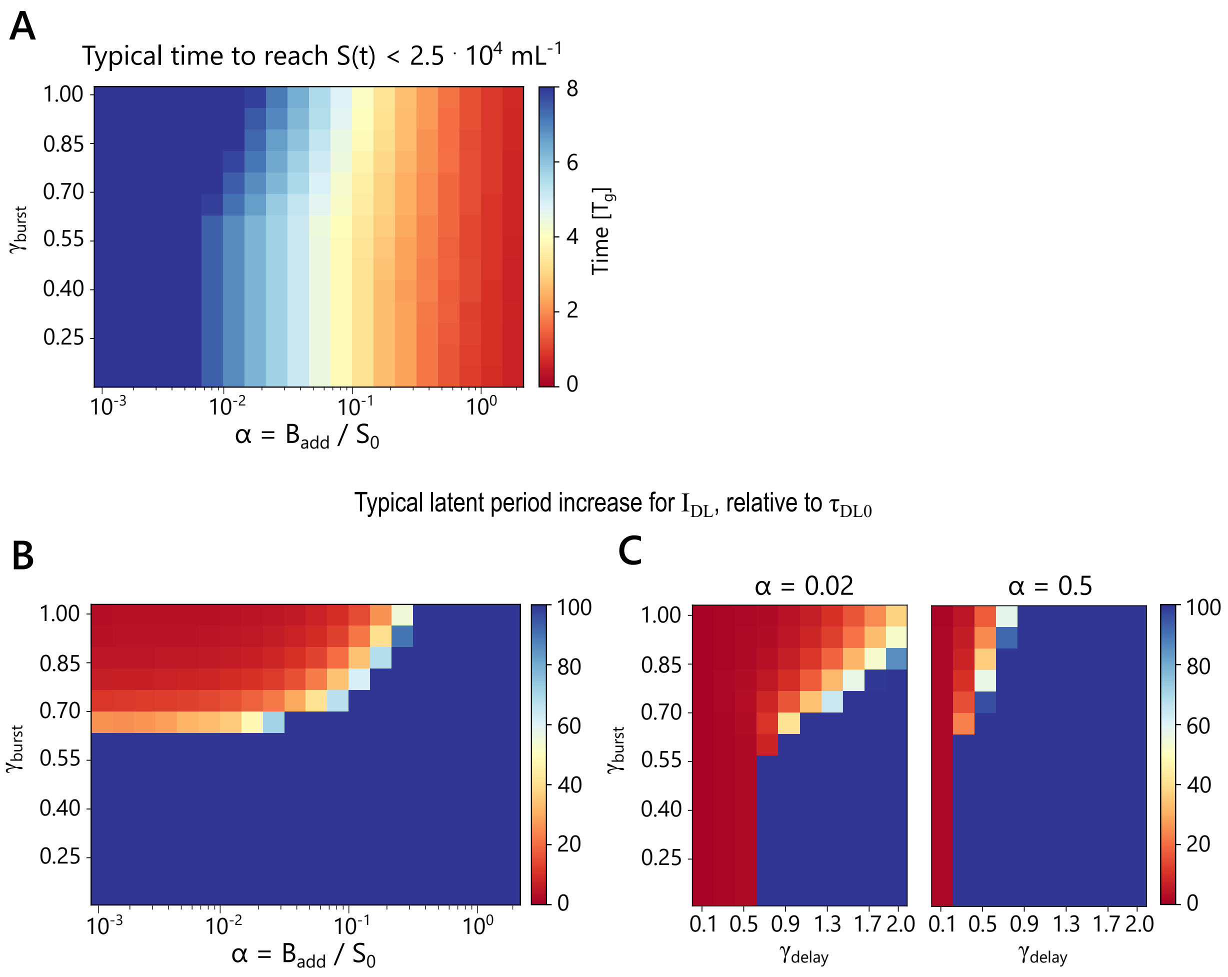


**SUPPLEMENTARY FIGURE** S4 Effective latent period increase of phage DL infected cells in the feast-famine system. A: Time index of the time point in a given cycle where $S$ typically drops below $S_{th}$ for different values of $\gamma_{burst}$ and $\alpha$. B: The typical latent period increase relative to $\tau_{DL_0}$ experienced by $I_{DL}$. Calculated as the weighted mean of $\tau_{DL}(S)$ over the time intervals where $\sum_{k=1}^{n} I_{DL}^{(k)} > 1$. C: Same as B, but for different values of $\gamma_{delay}$ and $\gamma_{burst}$ at $\alpha = 0.02$ and $\alpha = 0.5$.

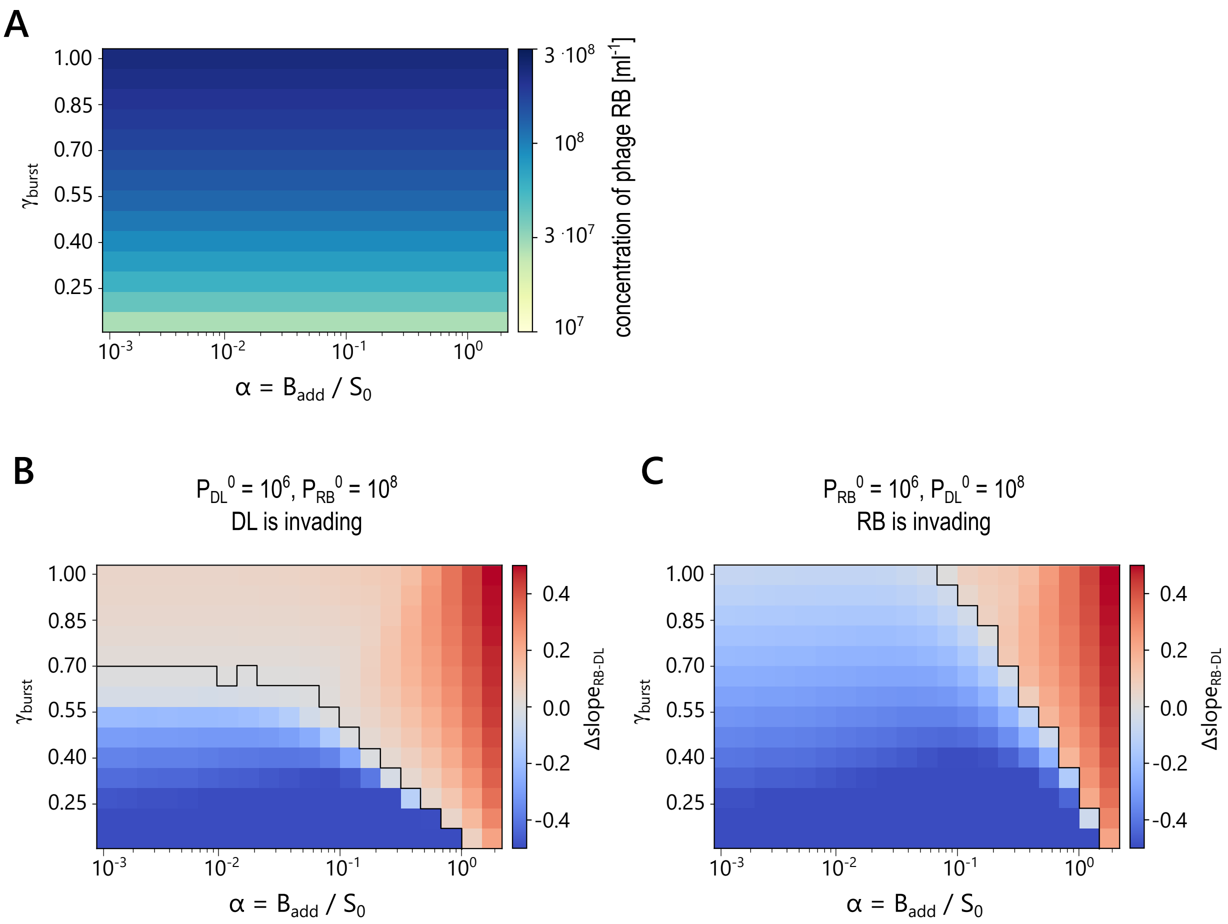


**SUPPLEMENTARY FIGURE** S5 Impact of initial conditions in the feast-famine system. A: The concentration of phage RB at the end of the first feast-famine cycle depends on $\gamma_{burst}$. B: The outcome of the feast-famine system in the tested parameter range remains relatively unchanged if the initial conditions are changed so that phage DL invades a system with dominant phage RB. C: If phage DL is set to be dominant at the start of the feast-famine simulation, the $\gamma_{burst}$ dependency of the outcome for low $\alpha$ is removed.

**A**

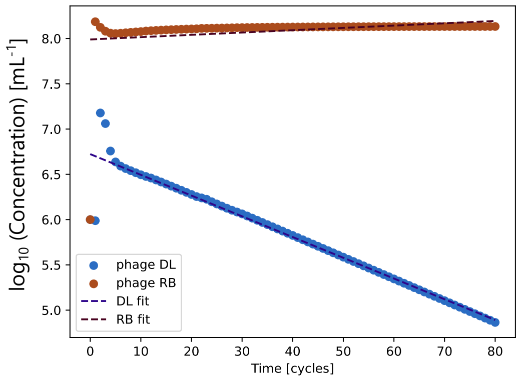


Extract $P(t_{end})$ for each cycle c

Fit linear regression model: $\log_{10}(P_{end}) = s_P \cdot c + b$

Final result is difference in slope:

$\Delta slope_{RB-DL} = s_{RB} - s_{DL}$

**B**

| Coexistence | Slow exclusion | Rapid exclusion |
|---|---|---|
| $\lvert\Delta slope\rvert < 0.0025$ | $0.0025 < \lvert\Delta slope\rvert < 0.07$ | $\lvert\Delta slope\rvert > 0.07$ |

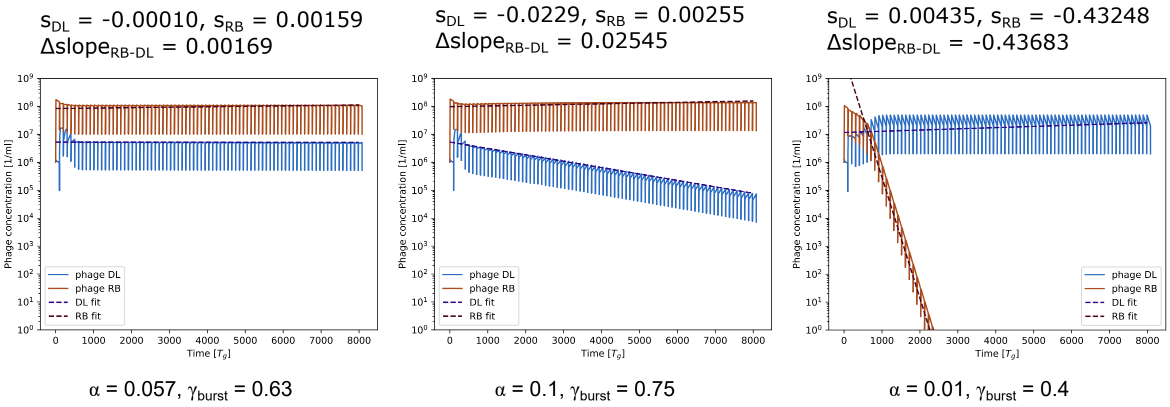


**SUPPLEMENTARY FIGURE** S6 Definition of the competition outcome value $\Delta slope$ in the feast-famine system. A: Explanation of linear regression approach. At the end of each feast-famine cycle, the value $P(t_{end})$ is extracted for both phages and stored in an array. At the end of the simulation, a linear model is fit to the $\log_{10}$ of the phage concentrations. The outcome of the competition is evaluated by the difference between the slopes of the two linear fits $\Delta slope_{RB-DL}$. As a result, a positive $\Delta slope$ outcome indicates that phage RB wins the competition and *vice versa*. B: We define two threshold values to differentiate between phage coexistence, slow exclusion, and rapid exclusion. We define slow exclusion as conditions in which both phages are present at significant concentrations ($> 10^2\ ml^{-1}$) over the simulated timescales (80 feast-famine cycles). This definition corresponds to $0.025 < \lvert\Delta slope\rvert < 0.07$.

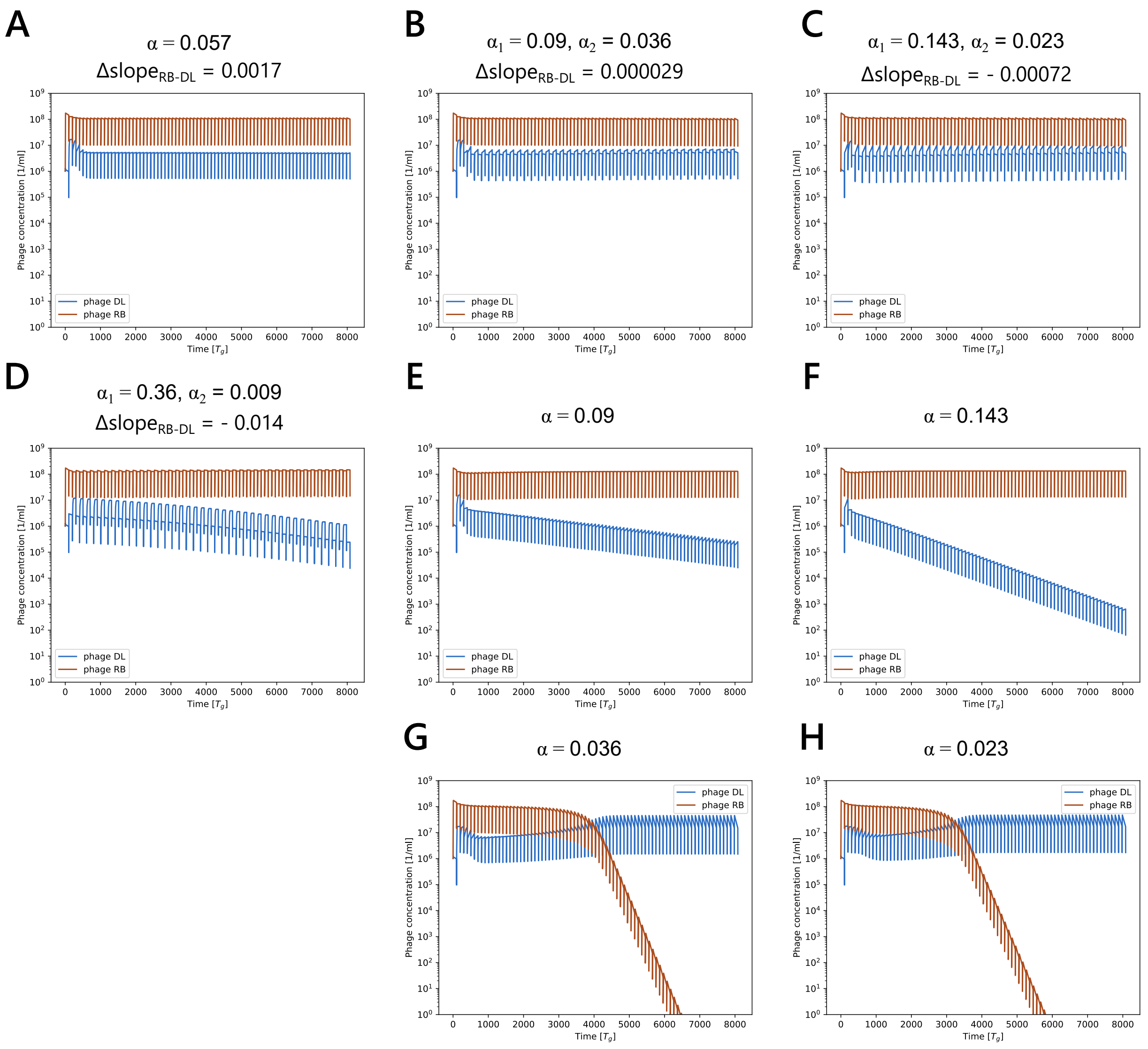


**SUPPLEMENTARY FIGURE** S7 Dynamics of phage DL and phage RB for simulations with alternating $\alpha$ values highlighted in Figure 4B with $\gamma_{burst} = 0.63$. A: Coexistence of the two phage types with a constant $\alpha = 0.057$. B, C: Coexistence of the two phage types with alternating $\alpha$ values with $\overline{\alpha} = 0.057$. D: Example of another combination of $\alpha_1$ and $\alpha_2$ with $\overline{\alpha} = 0.057$ which does not result in coexistence. E, G: Dynamics with constant $\alpha$ values from the pair in B (column wise). F, H: Dynamics with constant $\alpha$ values from the pair in C (column wise).

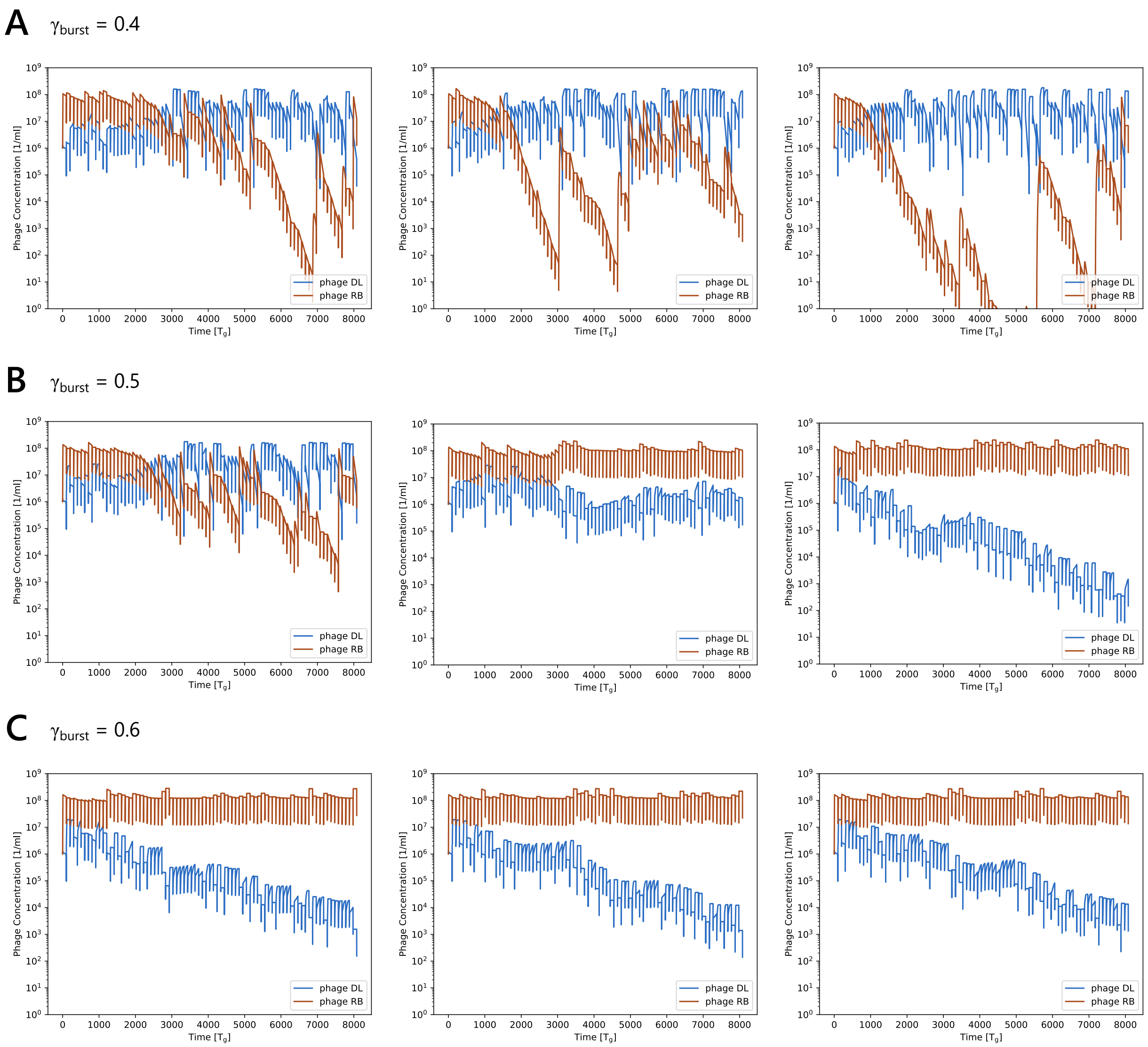


**SUPPLEMENTARY FIGURE** S8 Example dynamics of phage RB and phage DL under conditions of randomly selected $\alpha$ values at each reset event. A: $\gamma_{burst} = 0.3$, B: $\gamma_{burst} = 0.5$, C: $\gamma_{burst} = 0.6$.

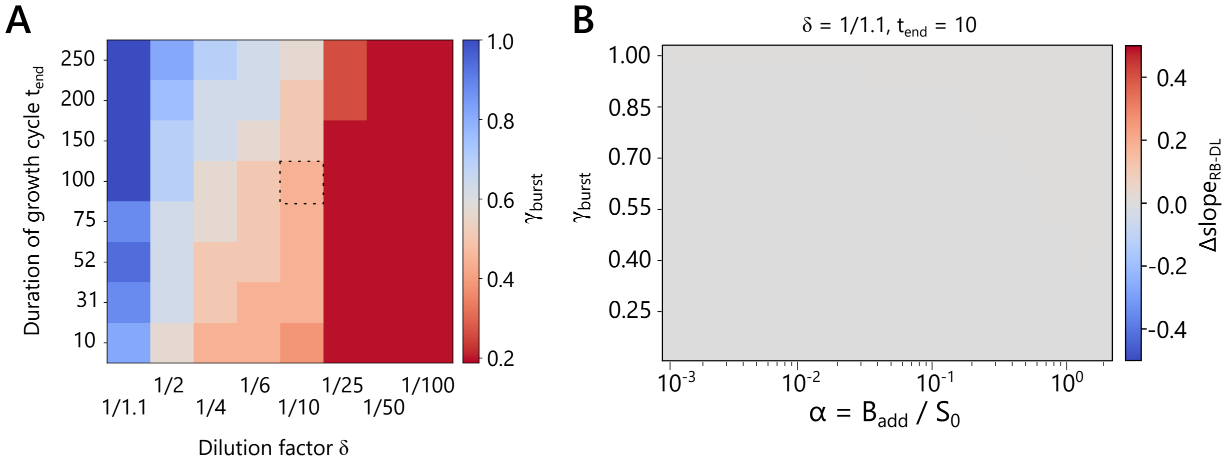


**SUPPLEMENTARY FIGURE** S9 Dependence of the outcome of the feast-famine system on the dilution factor $\delta$ and the duration of each growth cycle $t_{end}$. A: For each parameter combination 100 simulations were performed with randomly selected $\alpha$ values at each reset event, for 14 different values of $\gamma_{burst}$. The outcome is presented as the maximum $\gamma_{burst}$ parameter for phage RB that results in a negative median $\Delta slope_{RB-DL}$, i.e. the boundary at which the fitness of phage DL matches or exceeds the fitness of phage RB. Blue color indicates high fitness of phage DL, red color indicates low fitness of phage DL, respectively. The dotted square indicates the default conditions $\delta = 1/10,\ t_{end} = 100$ for all other figures. B: Outcome of phage competition for different values of $\gamma_{burst}$ and $\alpha$ at $\delta = 1/1.1 \approx 0.9$ and $t_{end} = 10$.

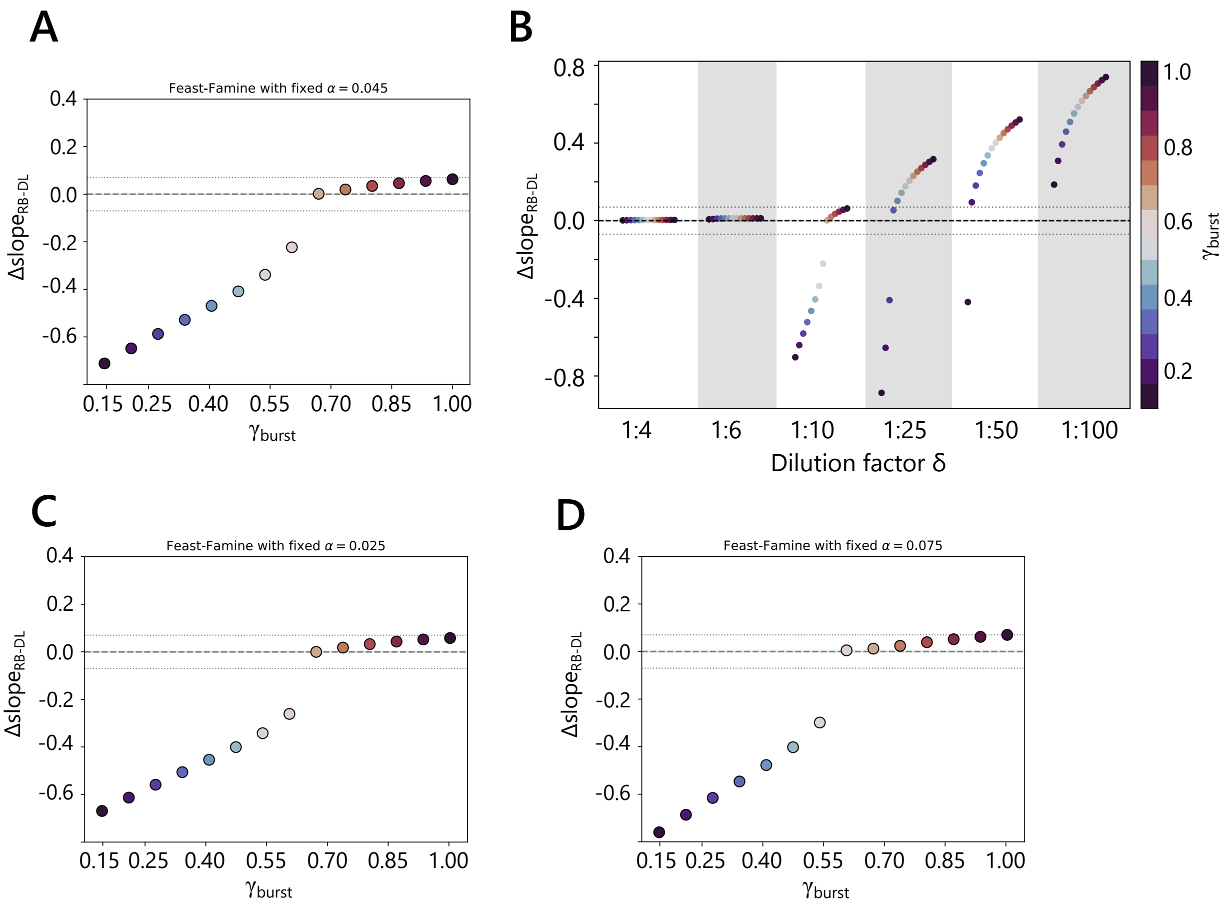


**SUPPLEMENTARY FIGURE** S10 Outcome of the feast-famine system with constant, average $\alpha$ instead of randomly fluctuating $\alpha$ value. A: In comparison to the fluctuating conditions in Figure 4C, phage DL fitness is higher in an environment with a constant $\alpha$ for the dilution factor $\delta = 1/10$. The geometric mean for the tested parameter range is $\overline{\alpha} = 0.045$. Dotted lines indicate the range defined as slow exclusion. B: Whether the fitness of phage DL is improved or diminished by the fluctuating conditions depends on the system dilution factor $\delta$. Compare with Figure 4D. For $\delta < 1/10$, any competitive advantage of phage DL is diminished by averaging of the environmental fluctuations, while phage DL fitness is enhanced by the averaging for $\delta \geq 1/10$ C, D: The averaged $\overline{\alpha}$ for 100 randomly selected values spans from $0.025 \leq \overline{\alpha} \leq 0.075$. Phage DL fitness is higher for the whole interval under conditions of a constant $\alpha$ compared to random fluctuations ($\delta = 1/10$).
